# Population dynamics with spatial structure and an Allee effect

**DOI:** 10.1101/2020.06.26.173153

**Authors:** Anudeep Surendran, Michael Plank, Matthew Simpson

## Abstract

Allee effects describe populations in which long-term survival is only possible if the population density is above some threshold level. A simple mathematical model of an Allee effect is one where initial densities below the threshold lead to population extinction, whereas initial densities above the threshold eventually asymptote to some positive carrying capacity density. Mean field models of population dynamics neglect spatial structure that can arise through short-range interactions, such as short-range competition and dispersal. The influence of such non mean-field effects has not been studied in the presence of an Allee effect. To address this we develop an individual-based model (IBM) that incorporates both short-range interactions and an Allee effect. To explore the role of spatial structure we derive a mathematically tractable continuum approximation of the IBM in terms of the dynamics of spatial moments. In the limit of long-range interactions where the mean-field approximation holds, our modelling framework accurately recovers the mean-field Allee threshold. We show that the Allee threshold is sensitive to spatial structure that mean-field models neglect. For example, we show that there are cases where the mean-field model predicts extinction but the population actually survives and vice versa. Through simulations we show that our new spatial moment dynamics model accurately captures the modified Allee threshold in the presence of spatial structure.

## 1 Introduction

Mathematical models of biological population dynamics are routinely built upon the classical logistic growth model where a population tends to a finite carrying capacity density for all positive initial densities [1–3]. While the logistic growth model is reasonable in some situations, there are other situations where the long-term survival of a population depends on the initial density, often called an Allee effect [4, 5]. A *strong Allee effect* is associated with a net negative growth rate at low densities leading to population extinction below the Allee threshold density. In contrast, a net positive growth rate at higher densities leads to the survival of the population when the initial density is greater than the Allee threshold [6–8]. Another type of Allee effect, known as the *weak Allee effect*, describes population growth with a reduced but positive growth rate at low densities [5]. Unlike the strong Allee effect, a weak Allee effect does not exhibit any threshold density due to the net positive growth rate. In this study, we focus on the strong Allee effect since we are interested in exploring threshold effects and factors that influence the Allee threshold. Initial evidence for the Allee effect came from ecological systems for various plant and animal populations [6, 9–15], whereas more recent studies suggest a role for the Allee effect in populations of biological cells [16–21].

Most mathematical models of Allee population dynamics invoke a mean-field assumption [2, 6, 22, 23] where, either implicitly or explicitly, interactions between individuals are assumed to occur in proportional to the average density. Such models neglect spatial correlations between the locations of individuals [24]. When short-range interactions are present, such as short-range competition, the mean-field approximation can become inaccurate [25–28]. Short-range interactions can lead to the development of spatial structure that can affect the overall population dynamics [29–31]. Spatial structure in biological populations includes both clustering and segregation [32–37]. Stochastic individual-based models (IBM) offer a straightforward means of exploring population dynamics without invoking a mean-field approximation [38, 39]. However, IBM approaches are computationally prohibitive for large populations and provide limited mathematical insight into the population dynamics, for example how particular biological mechanisms affect the carrying capacity or the Allee threshold [40].

A continuum approximation of the IBM in terms of the dynamics of spatial moments is a useful way to study how short-range interactions and spatial structure can influence population dynamics [41, 42]. Law et al. developed a spatial moment model, called the *spatial logistic model*, that quantifies the impact of spatial structure on classical logistic growth dynamics [29]. This work shows that spatial structure has a strong impact on the carrying capacity density. More recently, the spatial logistic model has been extended to consider other relevant mechanisms including interspecies and intraspecies interactions, neighbour-dependent motility bias, predator-prey dynamics and chase-escape interactions [43–46].

In this work, we present an IBM and a novel spatial moment dynamics approximation that incorporates a strong Allee effect. This model is a generalisation of the spatial logistic model [29]. Localised density-dependent interactions, such as short-range competition, short-range cooperation and short-range offspring dispersal are incorporated. In the limit of large scale interactions where the mean-field approximation is valid, both the IBM and the spatial moment model are consistent with the classical mean-field Allee growth model. In contrast, when spatial structure is present, we find that the Allee threshold density can be very sensitive to the spatial structure. For example, under a combination of short-range competition and short-range dispersal, all initial densities lead to population extinction, whereas the classical mean-field model predicts that the population will survive. In contrast, the new spatial moment approximation gives an accurate prediction of the long-time outcome even when strong spatial structure is present.

In this work we broadly consider two different types of results. In the first set we focus on population-level outcomes with regard to whether a population eventually becomes extinct or whether it survives. We study these problems using a classical mean-field model, a new spatial moment dynamics model as well as using repeated, identically-prepared stochastic simulations. For these results we are very careful to select initial conditions in the IBM simulations so that the vast majority of the repeated simulations lead to the same long-time outcome. For example, we consider parameter choices and initial conditions where more than ¿99% of identically-prepared IBM simulations all lead to either long-term extinction or long-term survival. The extremely small proportion of outliers are then excluded from the calculation of ensemble data. In the second set of results we focus on repeated IBM simulations in situations where stochastic effects can lead to either long-term survival or long-term extinction. We characterise this transition in terms of a survival probability, and we note that neither the classical mean-field or the new spatial moment dynamics model can make such predictions because they describe population-level outcomes only.

## 2 Individual-based model

The IBM describes the dynamics of *N* (*t*) individuals, initially distributed randomly on a continuous two-dimensional domain of size *L* × *L*. The location of the *n*^th^ individual is **x**_*n*_ ∈ ℝ^2^, and periodic boundary conditions are imposed. Individuals undergo birth, death and movement events, with event rates influenced by interactions between individuals. The IBM is developed for a spatially homogeneous, translationally-invariant environment, where the probability of finding an individual in a small region, averaged over multiple realisations of the IBM, is independent of the location of that region [43,46]. Hence the model is relevant to populations that do not involve macroscopic gradients in the density of individuals [47].

Competition between individuals influences the death rate, modelling increased mortality as a result of competition for limited resources. We use an interaction kernel, *ω*_*c*_(|***ξ***|), to describe the competition a particular reference individual experiences from another individual at a displacement, ***ξ***. We specify the competition kernel to be a function of separation distance, |***ξ***|,

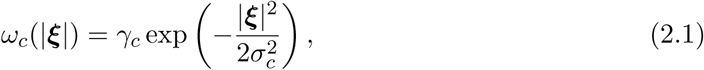

where *γ*_*c*_ > 0 and *σ*_*c*_ > 0 are the competition strength and range, respectively. Specifying the competition kernel to be Gaussian means that the impact of competition is a decreasing function of separation distance, |***ξ***|. We define a random variable, *X*_*n*_, that measures the neighbourhood density of the *n*^th^ individual weighted over by the competition kernel as,

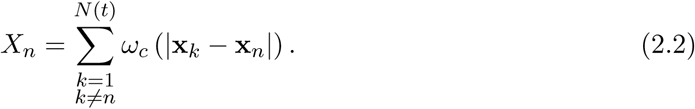

We consider the death rate of the *n*^th^ individual to be some function of *X*_*n*_,

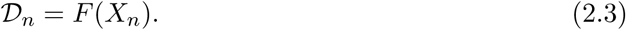

For a specific choice of *F* (*X*_*n*_), the key factors controlling how competition influences the death rate are *σ*_*c*_ and *γ*_*c*_. Figure 1 illustrates two scenarios for the simplest choice of *F* (*X*_*n*_) = *X*_*n*_. The arrangement of agents in Figure 1(a) and (c) are identical but we consider a long-range competition kernel (large *σ*_*c*_) in Figure 1(a)-(b) and a short-range competition kernel (small *σ*_*c*_) in Figure 1(c)-(d). Level curves of 𝒟_*n*_ are superimposed in Figure 1(a) and (c) and we see that the differences in the length-scale of interaction leads to very different local death rates. For example, when competition is long-range in Figure 1(a)-(b) the death rate for the relatively isolated green agent is 𝒟_*n*_ = 0.275 whereas when the competition is short-range in Figure 1(c)-(d) the death rate of the same agent is very different, 𝒟_*n*_ = 0. A similar set of results with a different choice of *F* (*X*_*n*_) in Appendix A shows similar results.

**Figure 1:**
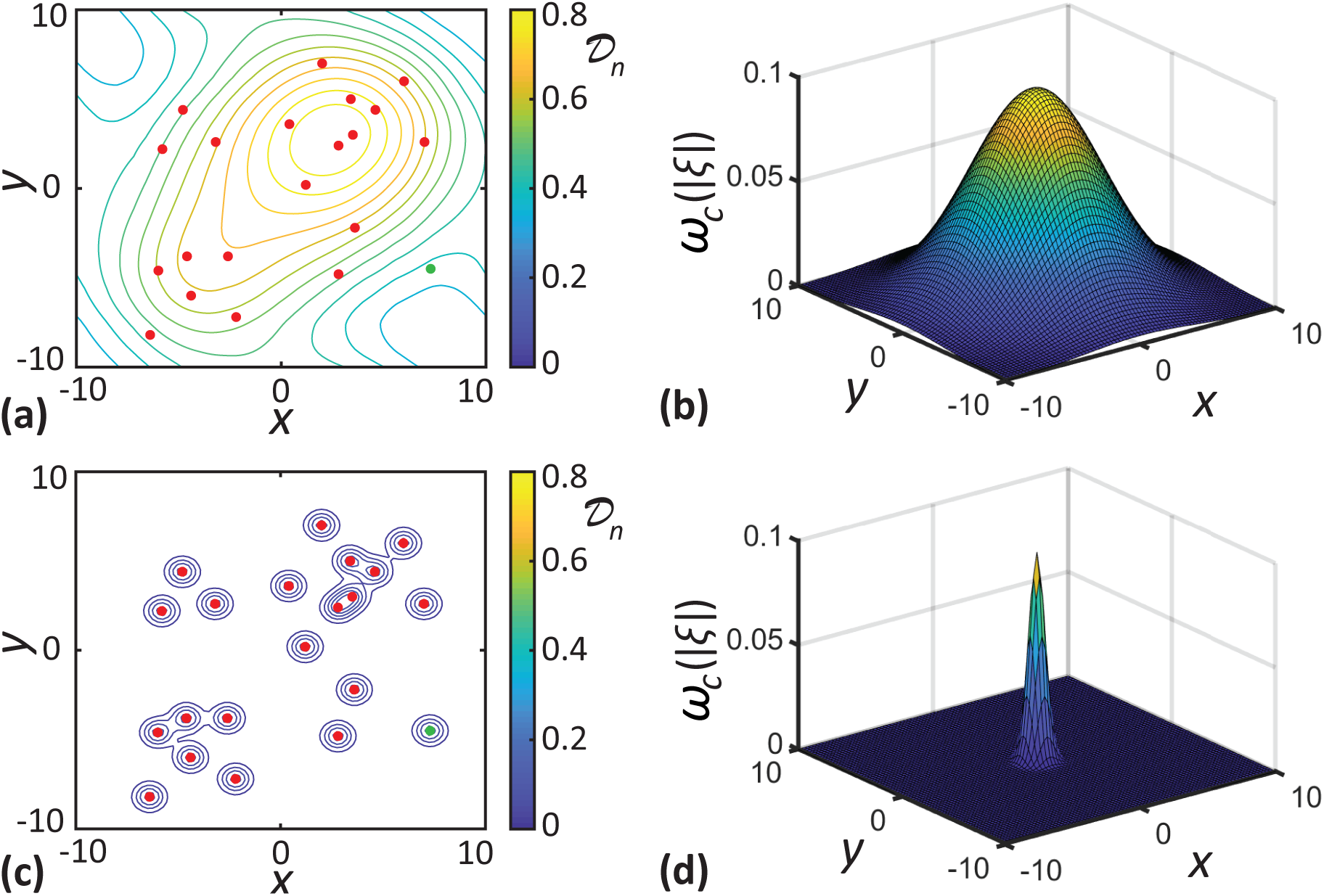
Visualising long- and short-range competition interactions. **a, c** locations of individuals (dots) superimposed with level curves of *D*_*n*_ for long- and short-range competition, respectively. **b, d** shows the long- (*σ*_*c*_ = 4.0) and short-range (*σ*_*c*_ = 0.5) competition kernels. Here, *F* (*X*_*n*_) = *X*_*n*_ and *γ*_*c*_ = 0.1.

We also consider a cooperative interaction between individuals that enhances the proliferation rate of individuals [19]. This is a model for sexual reproduction or some other mutualistic interaction in which reproductive fitness increases in the presence of near neighbours. Similar to the competition kernel, we define a cooperation kernel,

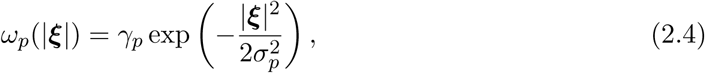

to account for the contribution of a neighbour at a displacement ***ξ*** to the reference individual’s proliferation rate. Here, *γ*_*p*_ > 0 and *σ*_*p*_ > 0 represent the strength and range of the interaction, respectively. As with competition, we define a random variable, *Y*_*n*_, that measures the neighbourhood density weighted by the cooperation kernel,

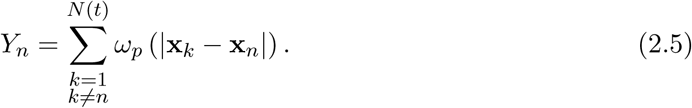

The proliferation rate of the *n*^th^ individual is taken to be some function of *Y*_*n*_,

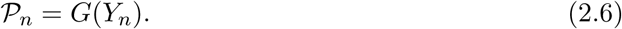

When an individual undergoes proliferation, a daughter agent is placed at a displacement sampled from a dispersal kernel, *µ*_*p*_(***ξ***) that we choose to be a bivariate normal with mean zero and standard deviation *σ*_*d*_.

For simplicity, we assume the movement rate is density-independent with a constant rate, *m*. An individual undergoing a motility event traverses a displacement, (|***ξ***| cos(*θ*), |***ξ***| sin(*θ*)) sampled from a movement kernel, *µ*_*m*_(***ξ***). The direction of movement, *θ* ∈ [0, 2*π*] is uniformly distributed. The distance moved, |***ξ***|, is sampled from a relatively narrow, truncated Gaussian distribution with mean, *µ*_*s*_, and standard deviation, *σ*_*s*_, where *σ*_*s*_ < *µ*_*s*_*/*4. To ensure |***ξ***| is positive, the Gaussian is truncated so that *µ*_*s*_ − 4*σ*_*s*_ < |***ξ***| < *µ*_*s*_ + 4*σ*_*s*_.

This IBM is an extension of the spatial stochastic logistic model [29], often simply called the spatial logistic model, which focuses on understanding the impact of short-range interactions and spatial structure on the classical logistic growth model [1]. In the spatial logistic model, the death rate is taken to be the sum of the competition from the neighbours and the proliferation rate is constant, so there is no cooperation. Furthermore, the spatial logistic growth model does not involve any agent motility. We recover the spatial logistic model as a particular case of our model when we set *F* (*X*_*n*_) = *X*_*n*_, *G*(*Y*_*n*_) = *p* and *m* = 0. In Figure 2(a)-(c), we summarise the dynamics of the spatial logistic model when the interactions are long-range and the mean-field approximation is valid. Under these conditions the death rate is a linearly increasing function of density and the proliferation rate is constant, as shown in Figure 2(a). This choice leads to an unstable steady state 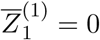 and a stable steady-state 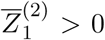 as shown in Figure 2(b). Hence the mean-field implies that all initial population densities will eventually tend to 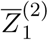 as *t* → ∞, as in Figure 2(c).

**Figure 2:**
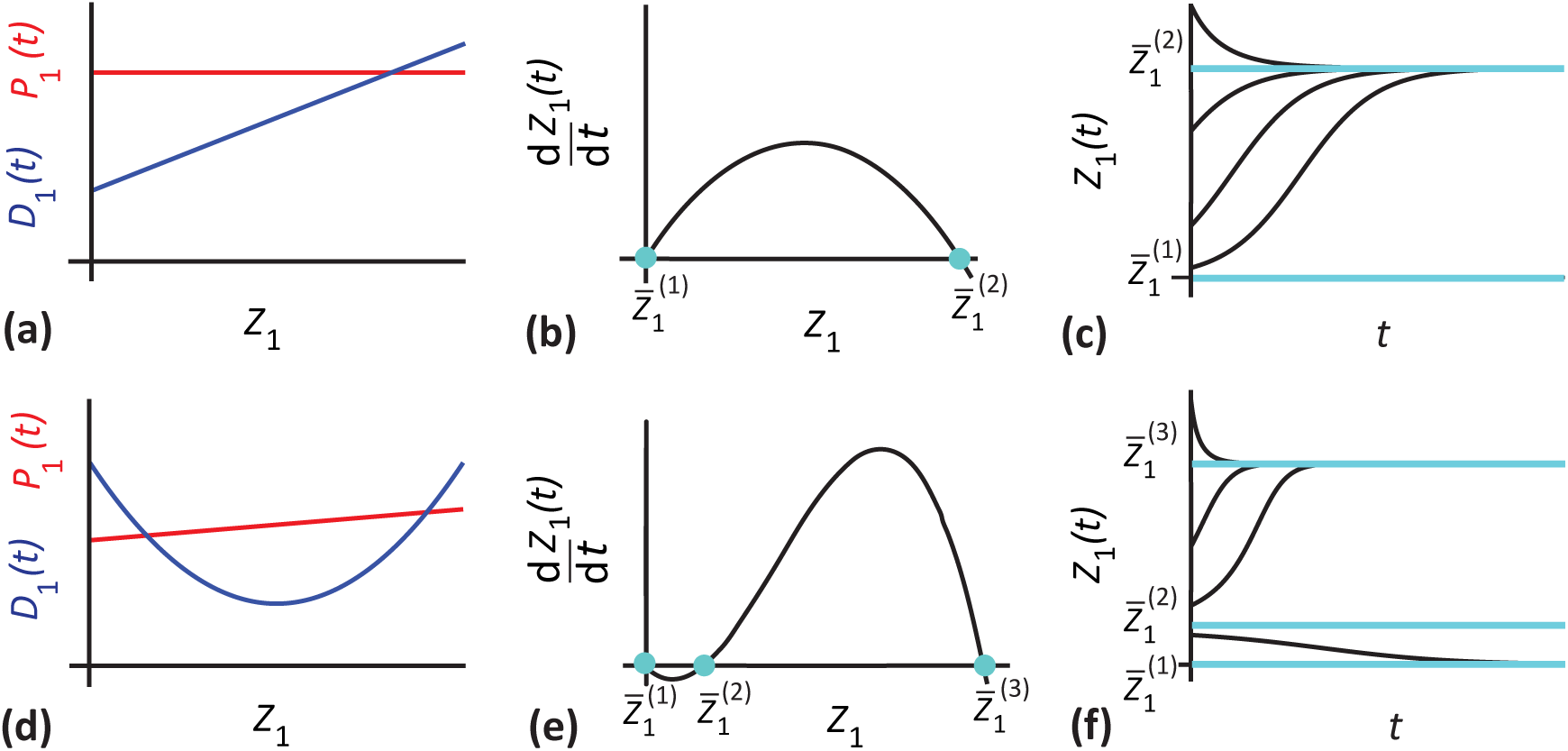
Comparison of the spatial logistic and Allee effect models under mean-field conditions. **a, d** proliferation (red) and death rates (blue) as functions of density. **b, e** density growth rate as a function of density. Equilibria highlighted with cyan dots. **c, f** dynamics for both models with the cyan lines indicating the equilibrium densities.

Our generalised IBM framework accommodates various non-linear functional forms for 𝒟_*n*_ and 𝒫_*n*_. In Figure 2(d) we choose 𝒟_*n*_ to be a concave up quadratic function and 𝒫_*n*_ to be linearly increasing function of density. This leads to three equilibrium densities: 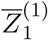,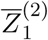 and 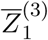, as in Figure 2(e). Here, 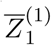 and 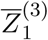 are stable equilibria, and 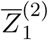 is an unstable equilibrium. The population dynamics here with the long-range interactions where the mean-field approximation is valid is shown in Figure 2(f). Here we see that populations with an initial density less than the Allee threshold, 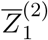, eventually go extinct. In contrast, any initial density greater than the Allee threshold, 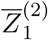, eventually tends to 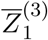. This is the simplest mean-field model of an Allee effect in which the net population growth rate is a cubic function of population density [3].

We simulate the IBM using the Gillespie algorithm [48] that is described in Section 2 of the Supplementary Material. The population dynamics arising from the IBM is analysed by considering the average density of individuals, *Z*_1_(*t*) = *N* (*t*)*/L*^2^. Information about the spatial configuration of the population can be studied in terms of the average density of pairs of individuals expressed as a pair correlation function, *C*(|***ξ***|, *t*) [31, 49, 50]. The pair-correlation function denotes the average density of pairs of individuals with separation distance |***ξ***|, at a time, *t*, normalised by the density of pairs in a population with the complete absence of spatial structure. Therefore, for a population without any spatial structure, *C*(|***ξ***|, *t*) = 1. When *C*(|***ξ***|, *t*) < 1, there are fewer pairs of individuals with a separation distance, |***ξ***|, than in a population without any spatial structure. We refer to this spatial configuration as being *segregated*. When *C*(|***ξ***|, *t*) > 1, we have more pairs of individuals with a separation distance, |***ξ***|, than we would have in a population without any spatial structure and this spatial configuration is referred to as being *clustered*.

## 3 Spatial moment dynamics

In this section, we construct a continuum approximation of the IBM in terms of the dynamics of spatial moments. The first spatial moment, *Z*_1_(*t*), for a point process of the kind considered in the IBM is defined as the average density of individuals [29, 43]. The second spatial moment, *Z*_2_(***ξ***, *t*), is the average density of pairs of individuals separated by a displacement of ***ξ***. The third spatial moment, *Z*_3_(***ξ, ξ*′**, *t*), is the average density of a triplet of individuals separated by displacements ***ξ*** and ***ξ***′, respectively. A formal definition of spatial moments is provided in Appendix C. It is possible to define higher-order moments similarly, but for the present study, we restrict our attention to the first three spatial moments [43, 51].

To derive dynamical equations for the evolution of the spatial moments, we need to find expressions for the continuum analogue of the discrete event rates given in Equation (2.3) and Equation (2.6). This can be achieved by finding the expected death and proliferation rates, 𝔼[*D*_*n*_] = 𝔼 [*F* (*X*_*n*_)] and 𝔼[*P*_*n*_] = 𝔼 [*G*(*Y*_*n*_)], respectively. To calculate these expected rates expand *F* (*X*_*n*_) and *G*(*Y*_*n*_) in a Taylor series about 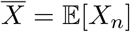 and 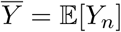. For the death rate we have

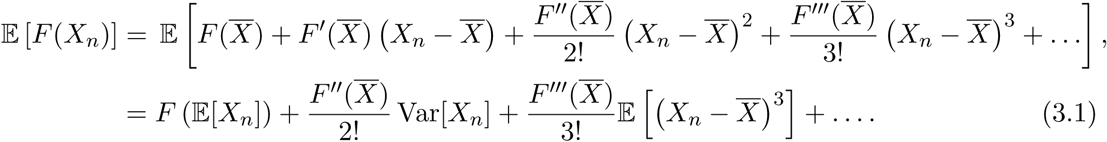

While our IBM can incorporate any choice of *F* (*X*_*n*_), the higher-order terms in the truncated Taylor series in Equation (3.1) are, in general, non-zero. The most straightforward choice of *F* (*X*_*n*_) to generate the Allee effect is a quadratic, and this choice has the additional benefit that third and higher derivatives vanish, so we have

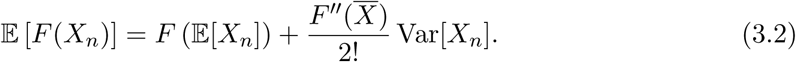

The computation of expected death rates reduces to calculating 𝔼[*X*_*n*_] and Var[*X*_*n*_], and substituting these into Equation (3.2). If we suppose the *L* × *L* domain is divided into *M* = *L*^2^*/*δ*A* subregions, each of area δ*A*, where these subregions are sufficiently small such that each subregion contains at most one individual, we have

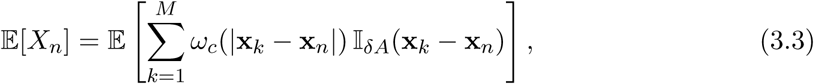

where, 𝕀_δ*A*_(**x**_*k*_ − **x**_*n*_) = 1, if an individual is present in a region of area δ*A* at a displacement **x**_*k*_ − **x**_*n*_, and 𝕀_δ*A*_(**x**_*k*_ − **x**_*n*_) = 0, otherwise. Using the property of the indicator function that 𝔼 [𝕀_δ*A*_(**x**_*k*_ − **x**_*n*_)] = ℙ [𝕀_δ*A*_(**x**_*k*_ − **x**_*n*_) = 1] [52], we have,

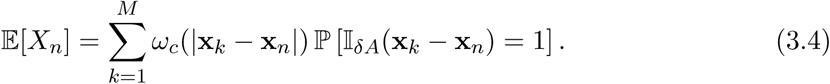

In the continuum limit, the right-hand side of Equation (3.4) is equivalent to multiplying the conditional probability of having an individual in a small window of size δ*A* at a displacement ***ξ*** from the reference individual, with the corresponding interaction kernel and integrating over all possible displacements as δ*A* → 0 [43]. The conditional probability for the presence of a neighbour individual is *Z*_2_(***ξ***, *t*) δ*A/Z*_1_(*t*). A derivation of the conditional probability is provided in Appendix D. Hence we have,

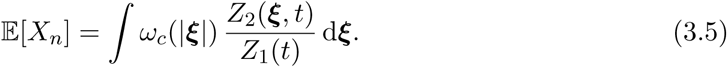

Note that the previous spatial moment models, including the spatial logistic model, assume that *F* (*X*_*n*_) is a linear function [29,30,40,43]. In that case, the death rate in Equation (3.2) depends only on 𝔼[*X*_*n*_]. But in the more general case where *F* (*X*_*n*_) is nonlinear, such as an Allee effect, we also require information about the variance. Therefore we compute Var[*X*_*n*_] in a similar fashion,

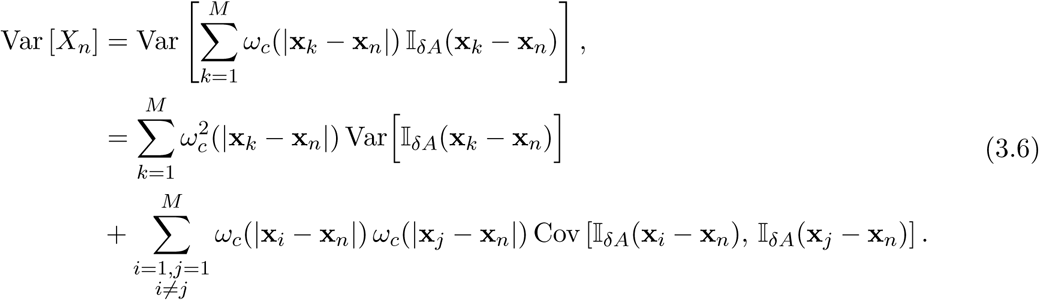

Following a similar procedure used in the computation of continuum analogue of 𝔼[*X*_*n*_] in Equation (3.5), we derive the expression for Var[*X*_*n*_] as,

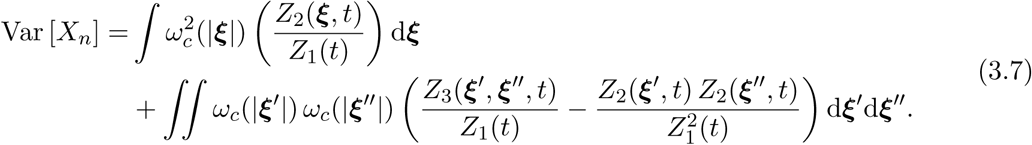

For brevity, we omit the intermediate steps involved in the derivation of Var[*X*_*n*_] here. These details are provided in Appendix E.

To make our spatial moment dynamics framework as general as possible, until this point we have made no assumptions about the choice of *F* (*X*_*n*_) and *G*(*X*_*n*_). From this point onwards we consider specific forms: 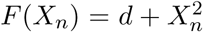 and *G*(*Y*_*n*_) = *p* + *Y*_*n*_. We make these choices because they are the simplest scenario that result in a strong Allee effect. To proceed, we compute the expected death rate of an individual, *D*_1_(*t*), by substituting the expressions for 𝔼[*X*_*n*_] and Var[*X*_*n*_] in Equation (3.2) to give,

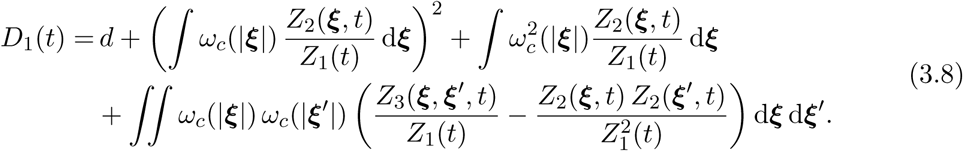

Similarly the expected proliferation rate for an individual is,

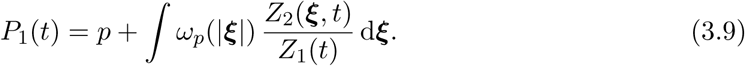

The dynamics of the first moment depend solely on the balance between proliferation and death. The movement of individuals does not result in a change in the population size. Hence the time evolution of the first spatial moment is given by,

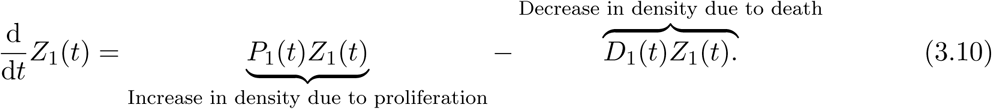

Note that the dynamics of the first moment depends on the second and third moments through Equations (3.8)-(3.9), and to solve the dynamics of the first moment, we need to specify the values of these higher-order moments.

Now we derive the dynamical equation for the density of pairs of individuals. For the derivation, we need to calculate the event rates of individuals while they are in a pair with another individual at a displacement, ***ξ***. The conditional probability of finding an individual at displacement ***ξ***′, given that a pair of individuals exist with separation displacement ***ξ*** is *Z*_3_(***ξ, ξ***′, *t*) δ*A/Z*_2_(***ξ***, *t*). The derivation of the expression for the conditional probability is given in Section 4 of the Supplementary Material. Using this expression for the conditional probability, and following the same procedures used to arrive at Equation (3.5) and Equation (3.7), we compute 𝔼[*X*_*n*_] and Var[*X*_*n*_] for an individual that forms a pair with another individual at a displacement ***ξ***. Hence, the expected death rate of an individual, conditional on the presence of a neighbour at a displacement ***ξ***, is given by,

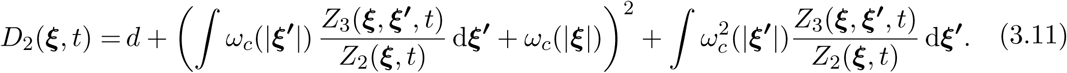

Note that the subscript in *D*_2_(***ξ***, *t*) indicates the fact that we are computing the expected rate for an individual that forms a pair with another individual at displacement ***ξ***. The additional factor of *ω*_*c*_(|***ξ***|) in the second term of Equation (3.11) accounts for the direct influence of the individual at displacement ***ξ***. Similarly, the expected proliferation rate of an individual, conditional on the presence of a neighbour at a displacement ***ξ*** is,

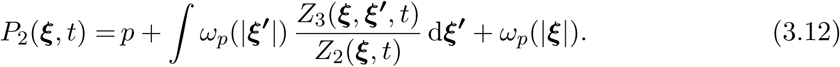

The time evolution of *Z*_2_(***ξ***, *t*) depends on the creation of new pairs and the loss of existing pairs. The schematic in Figure 3 illustrates possible ways in which movement, proliferation or death event leads to the creation or destruction of pairs of individuals separated by a displacement of ***ξ***. Figure 3(a) represents a pair of individuals at a separation displacement of ***ξ***. A movement of either individual destroys this pair, as does the death of either individual. Figure 3(b)-(c) demonstrates two different ways to generate a new pair separated by a displacement ***ξ***. When an individual among the pair separated by a displacement ***ξ*′** + ***ξ*** in Figure 3(b) moves or places a daughter agent a displacement ***ξ***, a new pair is formed at a displacement ***ξ***. In this case, the movement and proliferation occurs with rates *µ*_*m*_(***ξ*′**) *m* and *µ*_*p*_(***ξ*′**)*P*_2_(***ξ*′** +***ξ***, *t*), respectively. Another possibility for the creation of a pair with separation displacement ***ξ*** is when a single individual, as shown in Figure 3(c), places a daughter agent over a displacement -***ξ***. The rate for this event is *µ*_*p*_(−***ξ***) *P*_1_(*t*). The dynamics of the second moment is obtained by combining these possibilities as,

**Figure 3:**
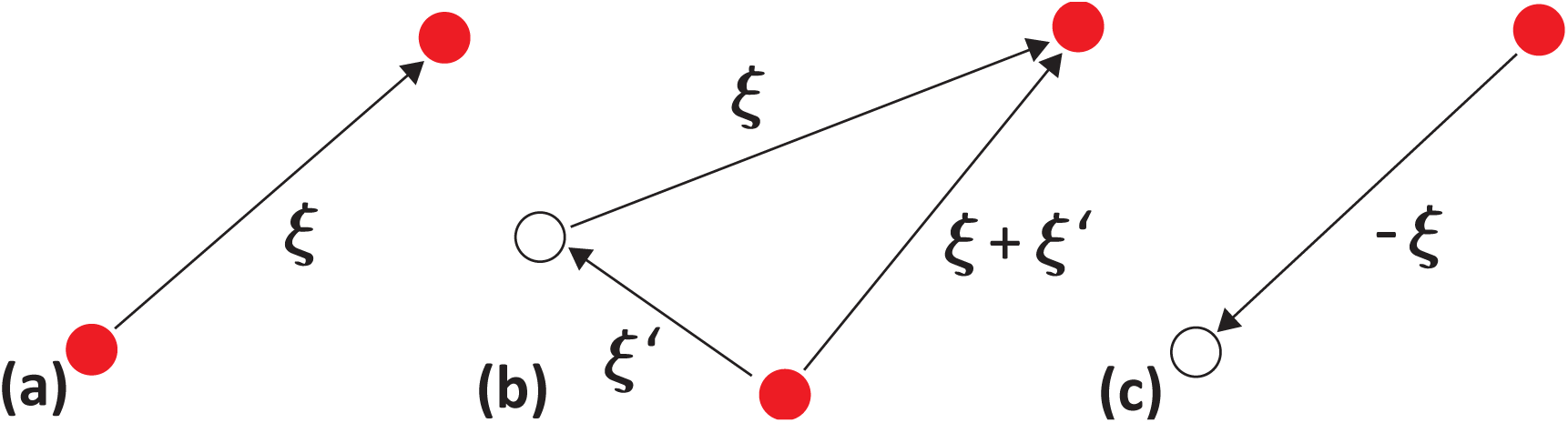
Possible events leading to a change in pair density. Red dots represent existing individuals and black open circles indicate potential locations of an individual after a movement or proliferation event. **a** A pair separated by a displacement ***ξ***. Movement or death of either individual destroys the pair. **b** A pair separated by a displacement ***ξ*** + ***ξ*′**. A movement or placement of a daughter over a displacement of ***ξ*′** creates a new pair separated by displacement ***ξ*. c** A single individual where the placement of a daughter at displacement **−*ξ*** creates a new pair with a displacement ***ξ***.

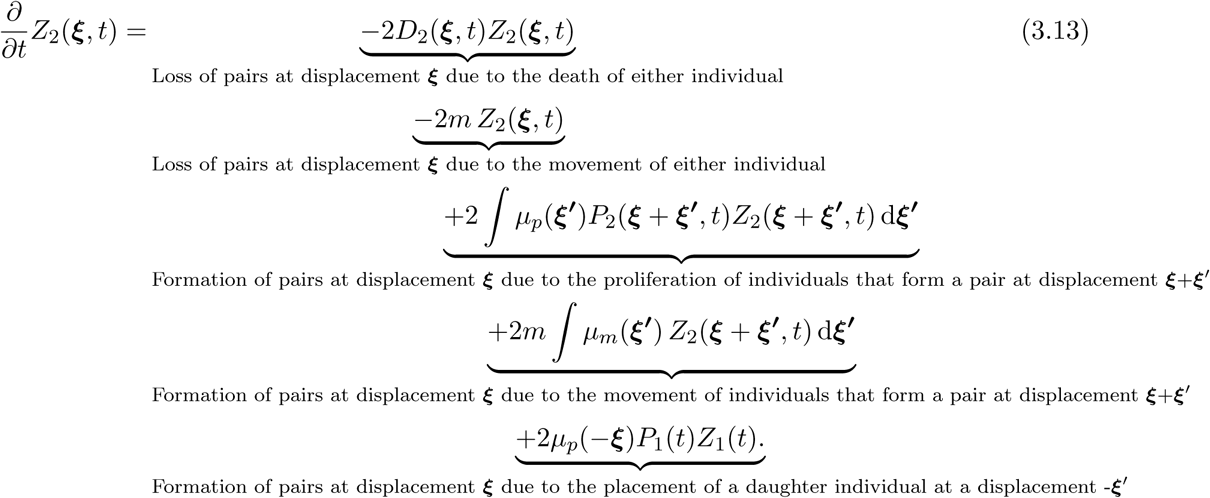

Since the event rates in Equations (3.11)-(3.12) depend on the third-order moment, *Z*_3_(***ξ, ξ***′, *t*), we need some expression for the third moment to solve the system, and we anticipate that the dynamics of the third moment will depend upon higher moments. To deal with this hierarchy of equations we use the Power-2 asymmetric moment closure approximation [29, 30]

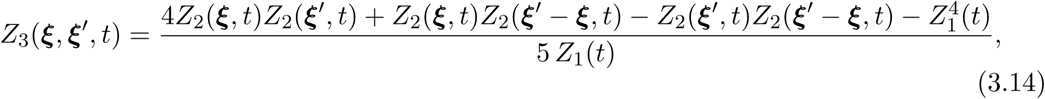

to approximately close the system in terms of the first and second moments only. Other closure approximations, such as the power-1 closure, the symmetric power-2 closure and the Kirkwood superposition approximation [29,53], are possible, and we compare the accuracy of these four different closure approximations in Appendix F. Details about the numerical methods involved in solving the dynamical equation for the second moment, Equation (3.13) is provided in Appendix G and MATLAB code to implement the algorithm is available on Github.

## 4 Mean-field dynamics

Under the classical mean-field approximation, interactions between individuals occur in proportion to the average density, and there is no spatial structure. These conditions correspond to having long-range interactions between individuals. Comparing the solutions of the classical mean-field model, IBM simulations and the solution of the new spatial moment dynamics model will provide insight into how spatial structure influences the population dynamics.

In terms of spatial moments, the mean-field implies that we have 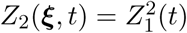 [30,31], which means that the expected death rate from Equation (3.8) simplifies to,

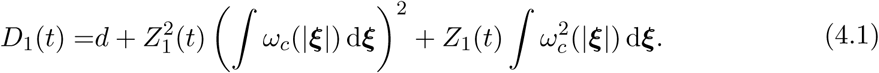

Similarly, the expected proliferation rate in Equation (3.9) simplifies to

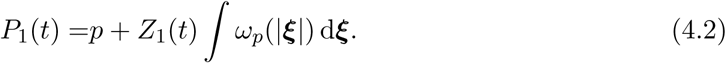

Since the interaction kernels have the property that 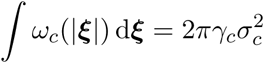 and 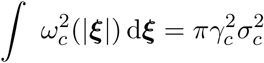, we substitute the mean-field death and proliferation rates in Equations (4.1)-(4.2) into Equation (3.10) to give

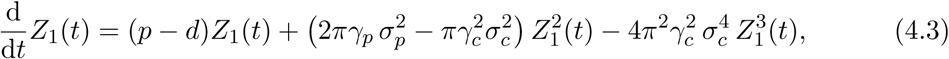

which leads to three equilibrium densities:

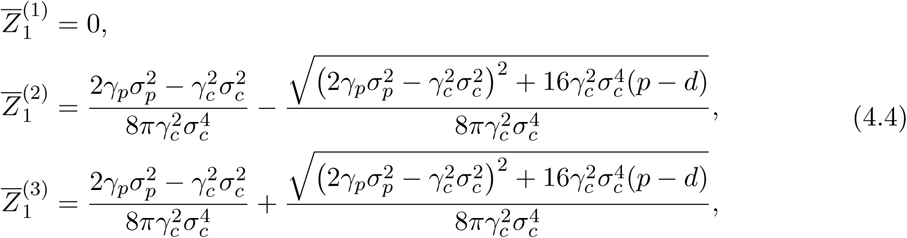

where 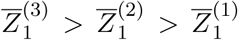 for the parameters we consider in this study. In this classical mean-field context, 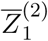 is the Allee threshold and 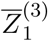 is the carrying capacity density. The equilibrium densities and dynamics associated with Equation (4.3) are depicted in Figure 2(e)-(f).

## 5 Results and Discussion

We now present IBM simulation results together with numerical solutions of both the spatial moment and the mean-field models to explore the influence of spatial structure on the population dynamics. In each case that we consider (Figures 4–7) we plot the time evolution of the average density of individuals from repeated, identically-prepared IBM simulations. Information about the spatial structure of the population is given in terms of the pair correlation function computed at the end of the simulation. Since the IBM is stochastic, there is a non-zero probability that any individual simulation will lead to extinction, regardless of whether the average outcome is that the population would survive. In the first set of results we present we take care to choose parameters and initial conditions such that at least 99% of the 1000 identically prepared simulations leads to the same long-term population-level outcome (i.e. extinction or survival) and any outliers, if present, are excluded from the calculation of ensemble averages. For each parameter combination, we consider three different initial conditions: 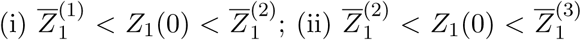; and 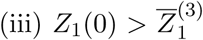. In the IBM simulations we control the initial density by choosing a different value of *N* (0).

**Figure 4:**
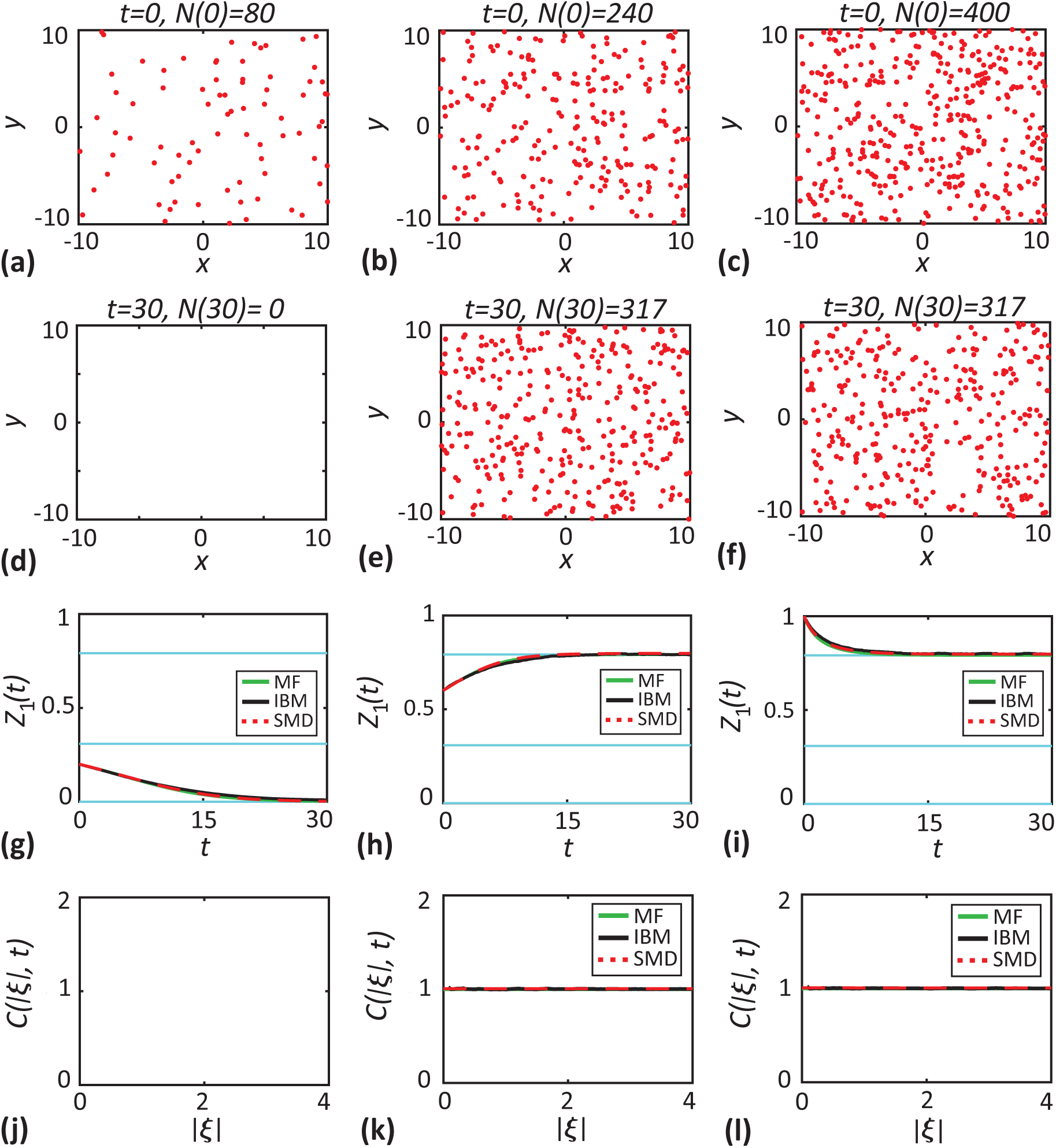
Long-range interactions and dispersal kernels (*σ*_*c*_ = *σ*_*p*_ = *σ*_*d*_ = 4.0) with weak interaction strengths (*γ*_*c*_ = 0.009 and *γ*_*p*_ = 0.009) lead to mean-field conditions. **a-c** initial locations of individuals (dots) for three different initial population sizes, *N* (0) = 80, 240 and 400, respectively. **d-f** location of individuals at *t* = 30. **g-i** density of individuals as a function of time. Black solid lines correspond to averaged results from 1000 identically-prepared IBM realisations, red dashed lines correspond to the solutions of the spatial moment dynamics model and green solid lines are the solution of the mean-field model. The cyan lines show the equilibrium densities. **j-l** pair-correlation function, *C*(|***ξ***|, 30). Parameter values are *d* = 0.4, *p* = 0.2, *m* = 0.1, *µ*_*s*_ = 0.4 and *σ*_*s*_ = 0.1.

As a starting point we consider a simple case with relatively weak long-range interactions so that the mean-field approximation is valid in Figure 4. Choosing long-range dispersion and competition kernels ensures that there is minimal correlation between the locations of agents in the simulations. As expected, results with 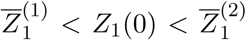 lead to extinction, and results with 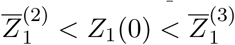 and 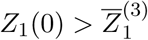 leads to the population eventually settling to the carrying capacity density by *t* = 30. The comparison of the averaged IBM results, the mean-field and spatial moment models in 4(g)-(i) shows that all three approaches are consistent and the eventual long-time population contains no spatial structure since the pair correlation function is unity in Figure 4(k)-(l). Note that we do not show the pair correlation function in Figure 4(j) since, in this case, the long-time result is that the population goes extinct.

Having verified that both the IBM and spatial moment dynamics model replicate solutions of the mean-field model for relatively weak long-range interaction and dispersal kernels, we now focus on short-range interactions that can lead to spatial structure.

### 5.1 Short-range competition reduces the Allee threshold

Results in Figure 5 are presented in the same format as in Figure 4 except that we consider short-range competition by setting *σ*_*c*_ = 0.5 without changing the cooperation or dispersal kernels.

**Figure 5:**
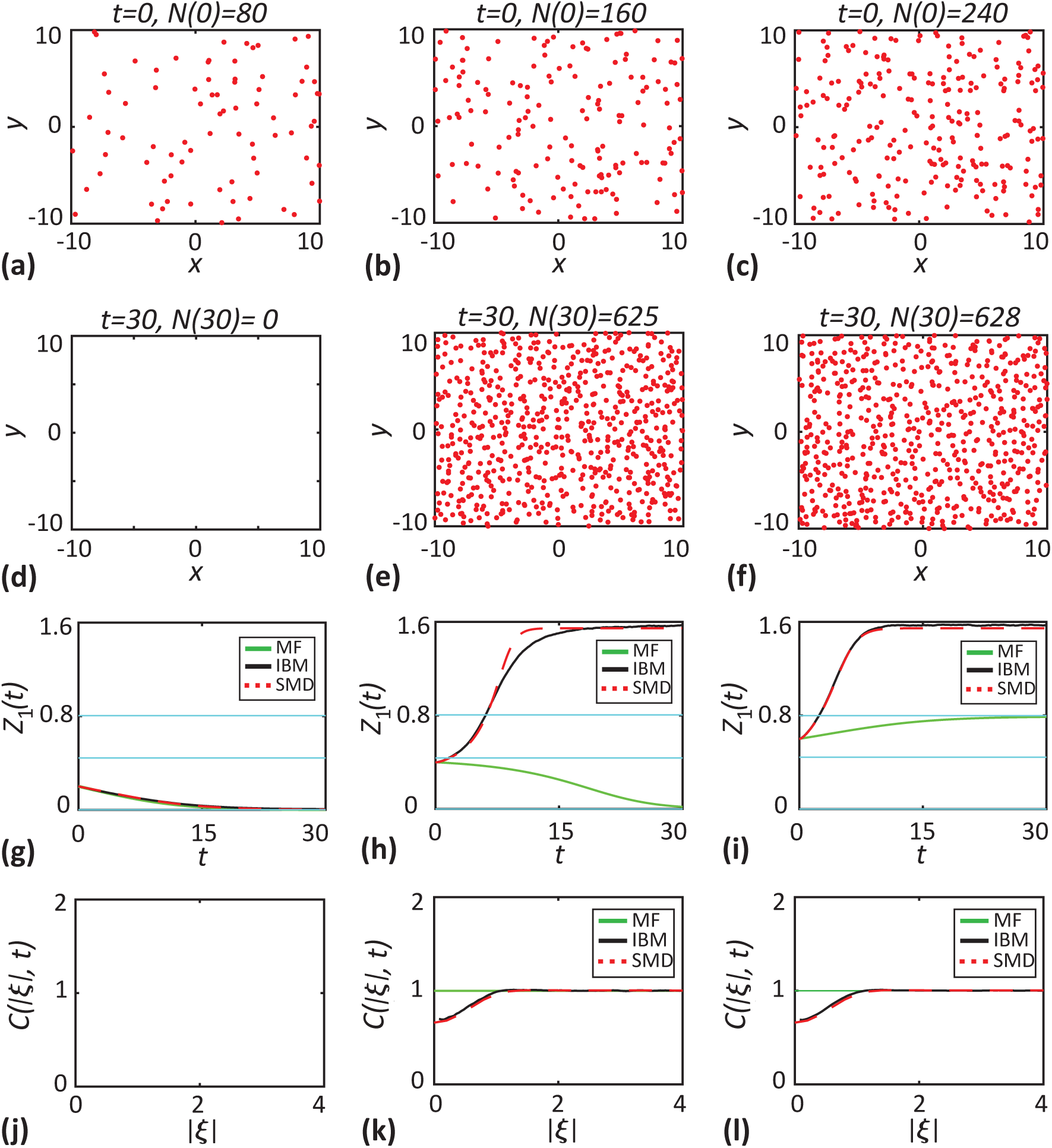
Short-range competition (*σ*_*c*_ = 0.5 and *γ*_*c*_ = 0.448) reduces the Allee threshold. **a-c** initial locations of individuals (red) for three different population sizes, *N* (0) = 80, 160 and 240, respectively. **d-f** location of individuals at *t* = 30. **g-i** density of individuals as a function of time. Black solid lines correspond to the averaged results from 1000 identically-prepared IBM realisations, red dashed lines correspond to the solutions of the spatial moment dynamics model and green solid lines are the solution of the mean-field model. The cyan lines show the equilibrium densities. **j-l** pair-correlation function, *C*(|***ξ***|, 30). Parameter values are *d* = 0.4, *p* = 0.2, *m* = 0.1, *µ*_*s*_ = 0.4 and *σ*_*s*_ = 0.1.

Figure 5(a)-(c) shows the initial randomly-placed populations, *N* (0) = 80, 160 and 240, respectively. Results in the left-most column with 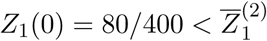 leads to extinction. Results in the central column with 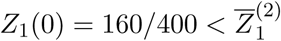 are very interesting because the initial density is below the classical mean-field Allee threshold and so standard models would predict extinction, yet we see that the population grows to reach a positive carrying capacity density. This difference is caused by the spatial structure, which we can see in Figure 5(k) is segregated at short distances. When competition is short range and the population is segregated, individuals in the IBM experience less competition that would be expected in a population without spatial structure under mean-field conditions. This decrease in competition means that the population increases despite the initial density being less than the mean-field Allee threshold.

Results in the right-most column in Figure 5 show that when the initial density is above the mean-field Allee threshold, 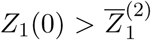, we see that the population increases to reach the same carrying capacity density is in Figure 5(h). Here we find that the carrying capacity density reached by the IBM is much higher than the mean-field carrying capacity density. This means that the classical mean-field model under-predicts the long-time density. In contrast, the spatial moment dynamics model leads to an accurate prediction of the averaged density from the IBM. This result, that the spatial structure can impact the carrying capacity density is consistent with previous observations for the spatial logistic model [29], but our observation that spatial structure changes the Allee threshold has not been reported previously.

### 5.2 Short-range competition and dispersal encourage population extinction

Results in Figure 6 consider both short-range competition and short-range dispersal by setting *σ*_*c*_ = *σ*_*d*_ = 0.5. The format of the results in Figure 6 is the same as in Figures 4-5. An additional set of results in Appendix H present some cases where we consider just short-range dispersal.

**Figure 6:**
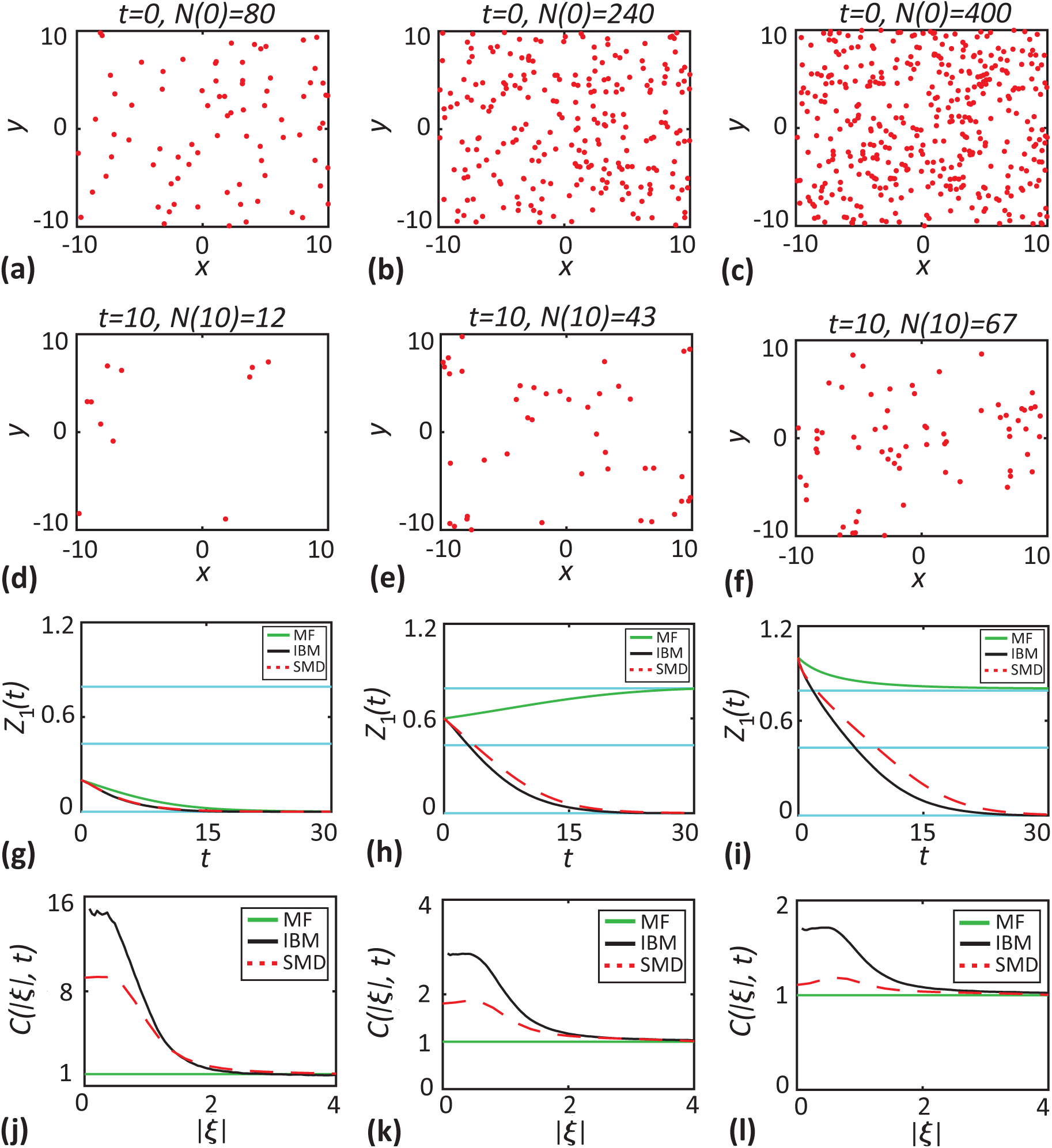
Short-range competition and short-range dispersal drive the population to extinction. In this case we consider short-range competition and dispersal (*σ*_*c*_ = *σ*_*d*_ = 0.5) with *γ*_*c*_ = 0.488. **a-c** initial locations of individuals (red) for three different population sizes, *N* (0) = 80, 240 and 400, respectively. **d-f** location of individuals at *t* = 30. **g-i** density of individuals as a function of time. Black solid lines correspond to the averaged results from 1000 identically-prepared IBM realisations, red dashed lines correspond to the solutions of spatial moment dynamics model and green solid lines are the solution of the mean-field model. The cyan lines show the equilibrium densities. **j-l** pair-correlation function, *C*(|***ξ***|, 30). Parameter values are *d* = 0.4, *p* = 0.2, *m* = 0.1, *µ*_*s*_ = 0.4 and *σ*_*s*_ = 0.1.

IBM simulations in Figure 6 show that short-range dispersal and competition leads to clustering and extinction. When the dispersal is short-range, daughter individuals are placed in close proximity to the parent individual, which leads to the formation of clusters. Short-range competition means that the competition within those clusters is strong, and significantly increases the local death rate of individuals. In the classical mean-field model we expect extinction to occur only when 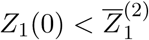 but our IBM results show that the population goes extinct when we set 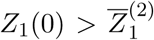 in Figure 6(h), and even more surprisingly we see that the IBM simulations lead to extinction even when 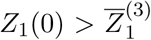 in Figure 6(i). This means that the spatial structure in this case leads the population to extinction. While the mean-field model completely fails to predict the observed extinction in the IBM, we find that the spatial moment model accurately reproduces the population dynamics of the IBM.

### 5.3 Short-range cooperation and intermediate-range dispersal promotes population growth

We now consider a further case where IBM simulations are qualitatively different from the classical mean-field model. Results in Figure 7 correspond to short-range cooperation (*σ*_*p*_ = 0.5) and intermediate-range dispersal (*σ*_*d*_ = 2.0). Some clustering in Figure 7(d)-(f) is evident, and this clustering is induced by the intermediate-range dispersal and leads to enhanced proliferation because of strong short-range cooperation. This case is very interesting because we observe population growth even when the initial density is below the classical mean-field Allee threshold in Figure 7(g). For all three choices of initial density in Figure 7, the population survives and eventually reaches a carrying capacity density that is greater than the carrying capacity density predicted by the classical mean-field model.

**Figure 7:**
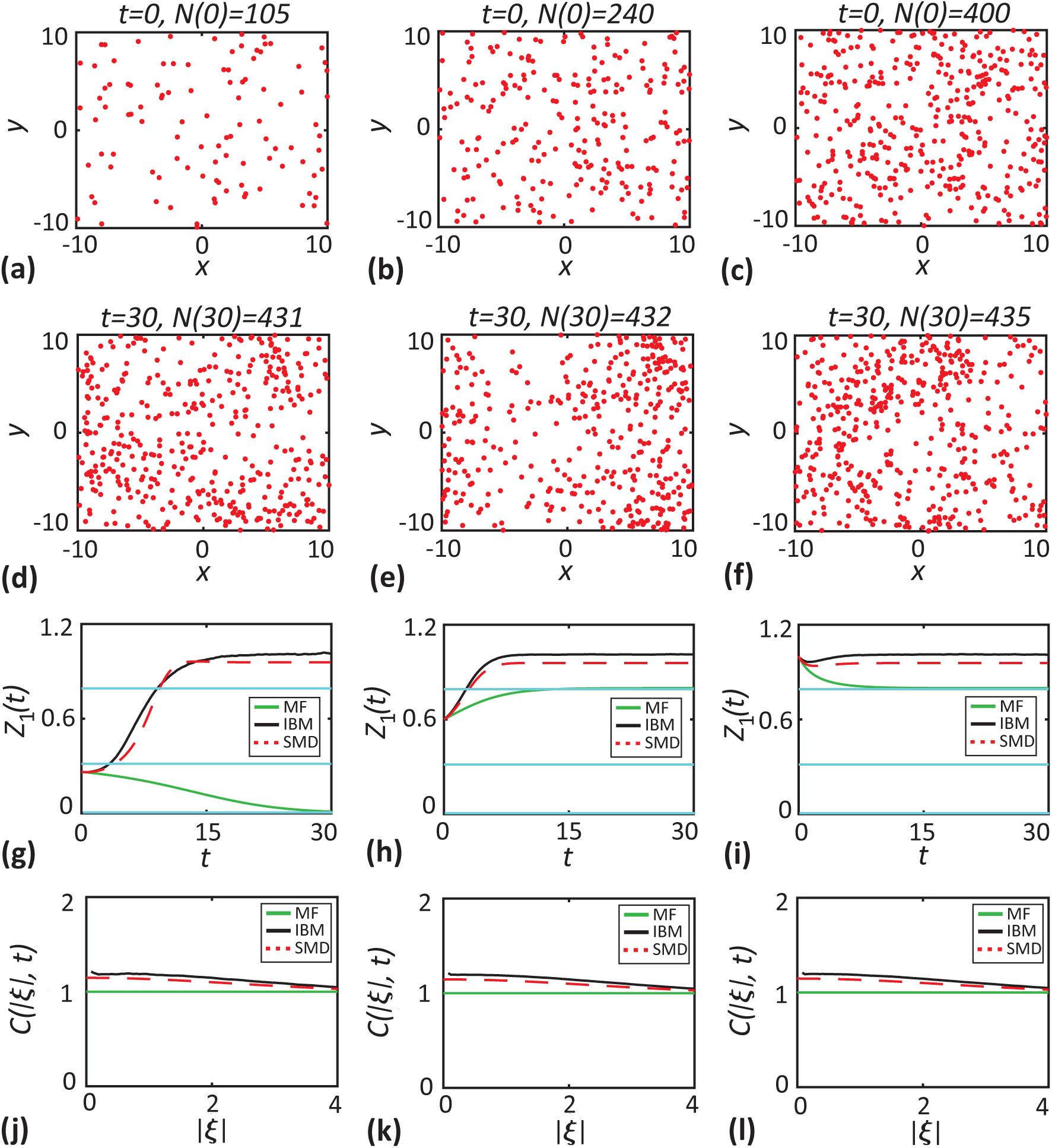
Short-range cooperation and dispersal promotes population growth. In this case we consider short-range cooperation *σ*_*p*_ = 0.5 and an intermediate-range dispersal *σ*_*d*_ = 2.0 with *γ*_*p*_ = 0.576. **a-c** the initial locations of individuals (red dots) for three different population sizes, *N* (0) = 105, 240 and 400. **d-f** show the location of individuals at *t* = 30. **g-i** show the density of individuals as a function of time. Black solid lines correspond to the averaged results from 1000 realisations of the IBM, red dashed lines correspond to the solutions of spatial moment dynamics and green solid lines correspond to the solution of the mean-field model. The cyan lines show the critical densities. **j-l** show the *C*(|***ξ***|, *t*) computed at *t* = 30 as a function of separation distance. Parameter values are *d* = 0.4, *p* = 0.2, *m* = 0.1, *µ*_*s*_ = 0.4 and *σ*_*s*_ = 0.1.

### 5.4 Survival probability

We conclude our results by using the IBM to estimate the survival probability, ℙ(survival), as a function of initial density in Figure 8. To calculate ℙ(survival), we perform a large number of identically-prepared realisations over a sufficiently long period of time, *t* = 100. From these simulations we record the fraction of realisations in which the population does not become extinct in this time interval. Results in Figure 8(a) show ℙ(survival) for a population with short-range competition. The mean-field Allee threshold for this choice of parameters is 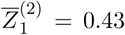. In contrast, we find that a certain proportion of populations with 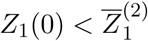 can survive. This difference between the classical mean-field result is due to the presence of spatial structure, and for this choice of parameters we have a segregated population, as illustrated in Figure 5. Another factor that influences the survival probability is the stochastic nature of the IBM. Even in the absence of spatial structure, the stochasticity can lead to a continuous transition of ℙ(survival) from 0 to 1 in the IBM, but possibly quite narrow and centred around the mean-field model Allee threshold. The spatial structure can shift the transition left or right (depending on whether it makes survival more or less likely) and potentially broaden the transition curve. Results in Figure 8(b) show ℙ(survival) for a population with short-range cooperation and intermediate-range dispersal where the classical mean-field Allee threshold is 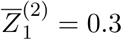. Again we see that populations with an initial density below the Allee threshold can survive. This difference between the classical mean-field result is due to the presence of spatial structure, which in this case leads to clustering, as shown in Figure 7, and the stochasticity of the IBM.

**Figure 8:**
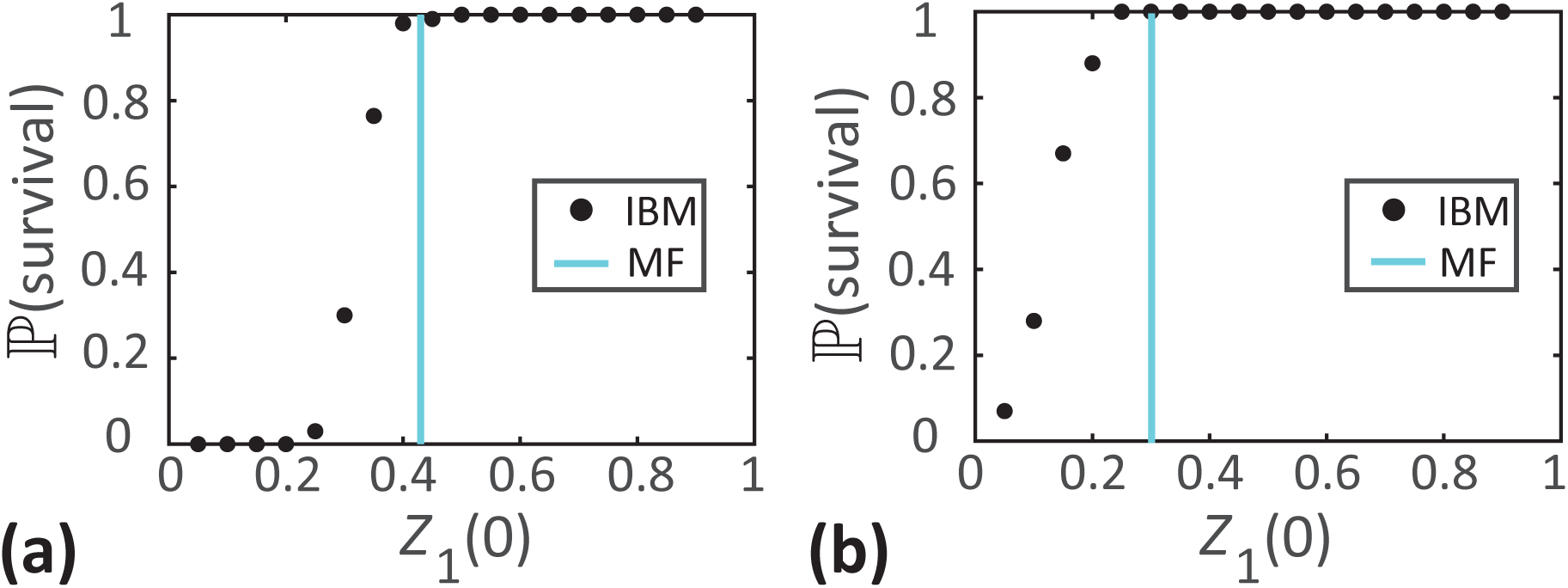
Survival probability as a function of *Z*_1_(0) for: **a** short-range competition (*σ*_*c*_ = 0.5, *γ*_*c*_ = 0.488), and **b** short-range cooperation and intermediate-range dispersal (*σ*_*p*_ = 0.5, *σ*_*d*_ = 2.0, *γ*_*p*_ = 0.576). Cyan lines show the classical mean-field Allee threshold and black dots show ℙ(survival) estimated from 100 identically-prepared IBM realisations. Other parameter values are *d* = 0.4, *p* = 0.2, *m* = 0.1, *µ*_*s*_ = 0.4 and *σ*_*s*_ = 0.1.

## 6 Conclusion and Outlook

In this study we consider an IBM of population dynamics that incorporates short-range interactions and spatial structure. The model construction allows us to incorporate a strong Allee effect, and in particular to explore the impact of spatial structure on the Allee threshold. Classical mathematical models of population dynamics that incorporate an Allee effect are based on making a mean-field approximation. This approximation implies that individuals interact in proportion to their average density and leads to the neglect of spatial structure, such as clustering and segregation.

We explore how short-range competition, short-range cooperation and short-range dispersal leads to spatial structure in a dynamic population and we focus on examining how this spatial structure influences the Allee threshold density. Overall, we find that the Allee threshold can be very sensitive to the presence of spatial structure to the point that classical mean-field predictions are invalid. For example, when we consider short-range dispersal and short-range competition we find that the population becomes extinct, despite the fact that the classical mean-field model predicts that the population will always survive when 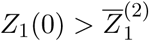. While our IBM results disagree with the classical mean-field prediction when the spatial structure is present, we also derive and solve a novel spatial moment dynamics model that is able to accurately capture how the Allee threshold depends upon spatial structure and we find that the spatial moment model reliably predicts population dynamics when spatial structure is present. Our results on the estimation of ℙ(survival) show that a certain proportion of populations seemed to have non-zero survival probability despite the fact that the initial population density is below the Allee threshold due to the presence of spatial structure. Note that the spatial moment model developed in this study is deterministic and hence cannot be used to estimate ℙ(survival). Instead, the spatial moment model predicts a threshold initial density above which the population survives. Overall our results show that the spatial moment model predicts this threshold more accurately than the mean-field model.

There are many potential avenues to extend the features in this study. For example, in this work we make the simplest possible assumption that agent movement is random and density-independent. This feature could be refined. For example, if the model was applied to study a population of biological cells, it might be more appropriate to consider a density-dependent movement rate and some directional bias where individuals are either attracted to or repelled from other agents in their neighbourhood [54,55]. In this study, we restrict our exploration to a simple population where all individuals are of the same type. Another interesting extension of the model would be to consider a multi-species where the total population consists of individuals from various distinct species [14,56]. We leave both these extensions for future consideration.

## Data Accessibility

Codes used for the simulation of the individual based model and numerical evaluation of spatial moment dynamics equation are available on Github.

## Acknowledgements

This study is supported by the Australian Research Council (DP170100474). MJP is partly supported by Te Pūnaha Matatini, a New Zealand Centre of Research Excellence.

## Appendices

### A Comparison of long-range and short-range competition

Here we present a second set of results comparing the impact of short-range and long-range competition on the death rate, 𝒟_*n*_, analogous to the results in Figure 1. These additional results, in Figure 9, are generated with precisely the same parameters and spatial arrangement of individuals as those in Figure 1, with the exception that here we consider a quadratic functional form for the death rate 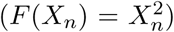. Comparing the results in Figure 1 with the additional results in Figure 9 indicates that the choice of 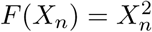, reduces the death rates of individuals.

**Figure 9:**
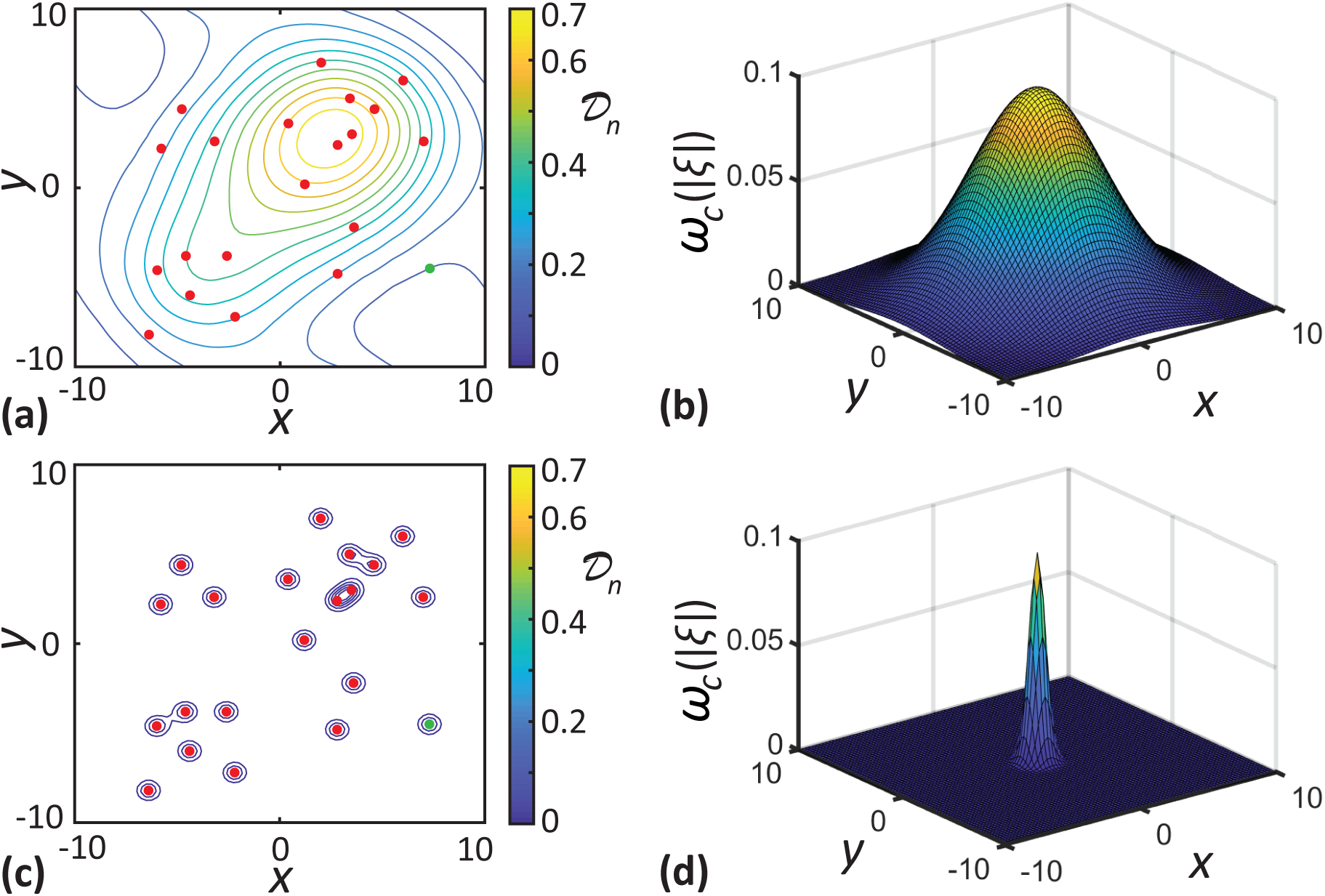
Visualisation of the impact of long-range and short-range neighbour-dependent interactions. Results in **a, c** show the location of individuals (red dots) superimposed with the level curves of 𝒟_*n*_ for long-range and short range-competition, respectively. Results in **b, d** show the long-range (*σ*_*c*_ = 4.0) and short-range (*σ*_*c*_ = 0.5) competition kernel, respectively, where these kernels are centred at the origin. For the computation of 𝒟_*n*_, we use 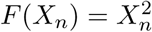 and *γ*_*c*_ = 0.1.

Similar to the results in Figure 1, the long-range competition leads to 𝒟_*n*_ being influenced by neighbours that are further apart, as shown in Figure 9(a)-(b). But the difference here is that overall we see a decrease in the death rate. For example, even though the death rate of the relatively isolated individual shown with the green dot is non-zero due to the longrange competition, the value of 𝒟_*n*_ = 0.076 here is lower than 𝒟_*n*_ = 0.275 computed using *F* (*X*_*n*_) = *X*_*n*_ in Figure 1. Under short-range competition, as shown in Figure 9(c)-(d), only the contribution from immediate neighbours is significant. As a result, the isolated individual (green dot), does not experience competition from its neighbours, leading to 𝒟_*n*_ = 0.

### B Numerical implementation of the individual-based model

Here we outline the numerical implementation of our individual-based model (IBM) using the Gillespie algorithm [48]. The codes used for the simulation of the IBM are available on Github. In each simulation, we initially populate the computational domain, of size *L* × *L*, with *N* (0) individuals placed at random. We use periodic boundary conditions along all boundaries. For each potential event, we compute the death and proliferation rates of individuals using Equation (2.3) and Equation (2.6). The movement of individuals is density independent, and the constant intrinsic movement rate is *m*. The sum of event rates of all individuals is given by,

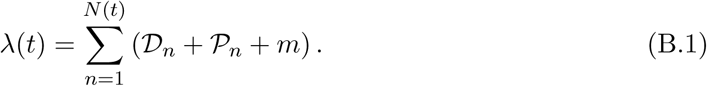

Each time the IBM is updated, one of the three possible events occurs, and the time interval between successive events is exponentially distributed with mean 1*/λ*(*t*). The probability for any of the events to occur is proportional to the rate of the corresponding event. For a proliferation event, an offspring is placed at a displacement sampled from the dispersal kernel, *µ*_*p*_(***ξ***), and the population size increases by one. For a death event, the population size reduces by one. For a movement event, an individual traverses a displacement that is sampled from the movement displacement kernel, *µ*_*m*_(***ξ***).

We compute the average density of individuals at a particular time by dividing the population size, *N* (*t*), by the area of the computational domain, *L*^2^. To compute the pair-correlation function, *C*(|***ξ***|, *t*), we consider a reference individual located at **x**_*n*_ and calculate all distances |***ξ***| = |**x**_*k*_ − **x**_*n*_|, associated with the other *N* (*t*) − 1 individuals [29, 40]. We repeat this process with each of the remaining individuals until every individual has acted as the reference individual. The pair-correlation function is calculated by enumerating the distances which fall into the interval, [|***ξ***| − δ|***ξ***|*/*2, |***ξ***| + δ|***ξ***|*/*2]. We normalise the bin count by a factor of 2*π*|***ξ***|δ|***ξ***|*N* (*t*)(*N* (*t*) − 1)*/L*^2^ to ensure that *C*(|***ξ***|, *t*) = 1 in the absence of spatial structure. The choice of bin width, δ|***ξ***|, is crucial in computing the pair-correlation function. When δ|***ξ***| is very small, we obtain a noise dominated *C*(|***ξ***|, *t*). In contrast, very large δ|***ξ***| leads to an overly smooth *C*(|***ξ***|, *t*) that fails to describe the effects of short-range interactions. In all our simulations, we use an intermediate value of δ|***ξ***| = 0.2 which helps us to avoid the two extremities.

### C Definition of spatial moments

Here, we provide a more formal mathematical definition for the spatial moments [24, 30, 43]. Let us suppose *D*_δ*A*_(**x**) ⊂ ℝ^2^ is a disc of area δ*A* centred at position **x** ∈ ℝ^2^ and the number of individuals in the region *D*_δ*A*_(**x**), at a time *t*, is denoted by the random variable *N* (*D*_δ*A*_(**x**), *t*). The first spatial moment, *Z*_1_(*t*), can be computed by dividing the population size of individuals by the area of the domain. Hence we have,

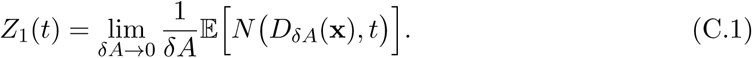

The second spatial moment, *Z*_2_(***ξ***, *t*), is the average density of pairs of individuals separated by a displacement ***ξ*** at time *t*. For a pair of individuals separated by a displacement ***ξ***, we have,

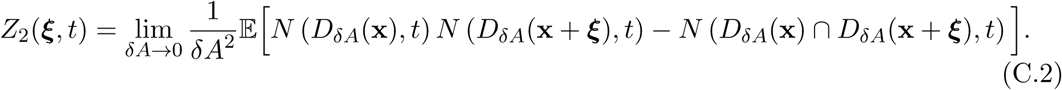

The second term in the expectation in Equation (C.2) is necessary to avoid counting self-pairs. For non-overlapping regions *D*_δ*A*_(**x**) and *D*_δ*A*_(**x** + ***ξ***), this term becomes zero as δ*A* → 0.

The third spatial moment is the density of triplets of individuals, and is similarly defined as,

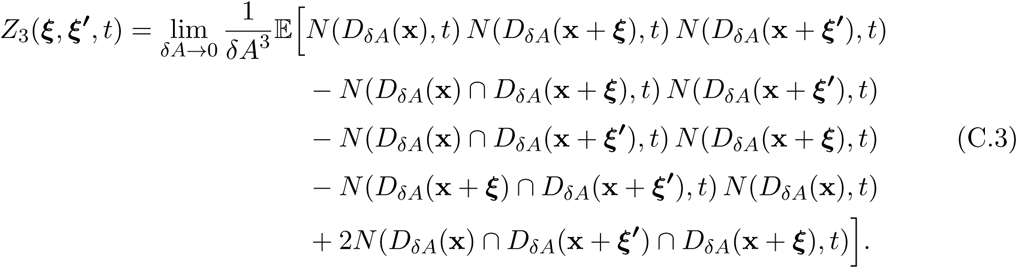

Again, the extra terms in Equation (C.3) are needed to avoid counting non-distinct triplets.

### D Conditional probabilities for the presence of individuals

In this section we derive expressions for the probabilities of finding individuals at specific displacements conditional on the presence of other individuals. In the limit, δ*A* → 0, the probability of having one individual in the region *D*_δ*A*_(**x**) is given by,

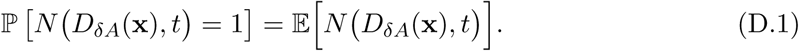

Now, the probability of having two individuals located in non overlapping regions *D*_δ*A*_(**x**) and *D*_δ*A*_(**x** + ***ξ***), respectively, is given by,

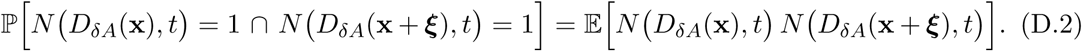

Similarly, the probability for having three individuals at non overlapping regions *D*_δ*A*_(**x**), *D*_δ*A*_(**x** + ***ξ***) and *D*_δ*A*_(**x** + ***ξ***′), respectively, is given by,

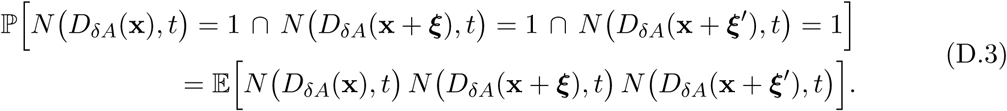

To compute the event rates, we need to find the probabilities of individuals being present at a given displacement, conditional on the presence of other individuals. To compute the conditional probabilities, we use the property that,

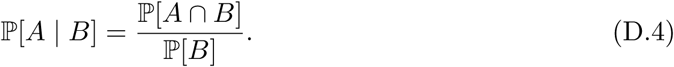

The conditional probability of finding an individual at a displacement **x** + ***ξ***, given that the reference individual is located at **x**, is,

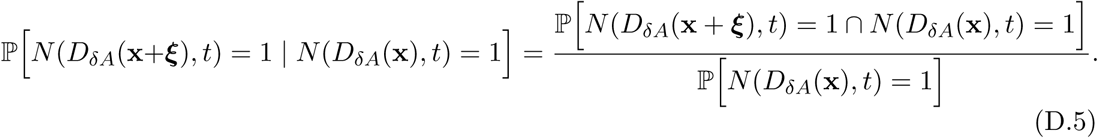

Using the definitions of probabilities in Equations (D.1)-(D.2) and the definitions of spatial moments in Equations (C.1)-(C.2), we rewrite the the numerator and denominator of Equation (D.5) as,

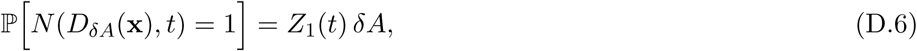

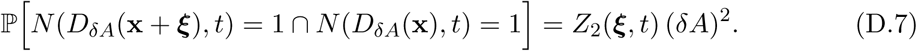

Hence the conditional probability of finding an individual at a displacement **x** + ***ξ*** from a reference individual at **x** is given by,

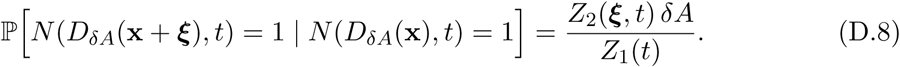

Similarly, we compute the conditional probability of finding an individual at a displacement **x** + ***ξ*′**, given that a pair of individuals where constituent individuals are located at **x** + ***ξ*** and **x**, respectively, as

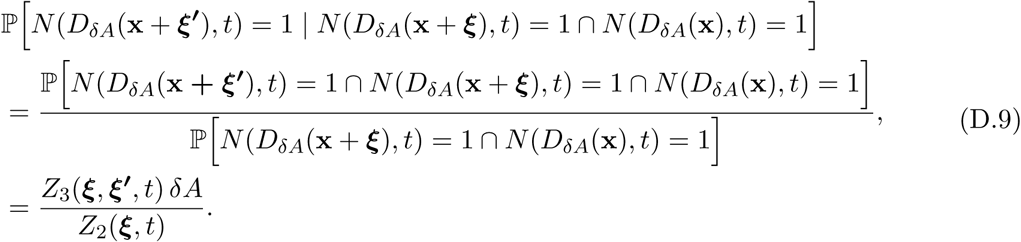

### E Computation of variance

Here we outline the steps involved in the derivation of expression for variance of the neighbourhood density in Equation (3.7). The fundamental definition of the variance is,

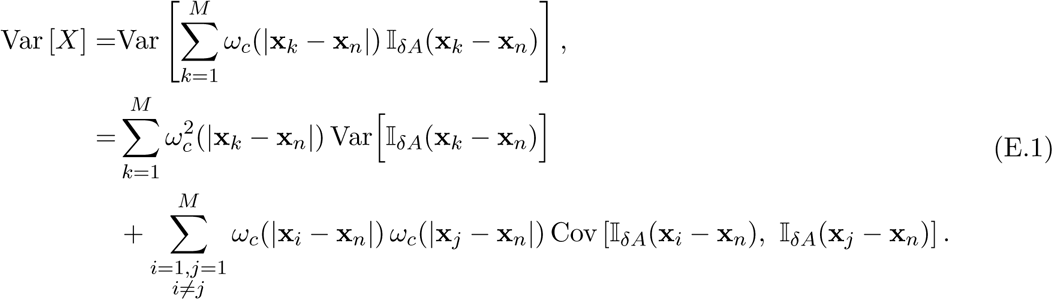

Now, using the properties of indicator function, [𝕀_δ*A*_(**x**_*k*_ − **x**_*n*_) = ℙ 𝕀_δ*A*_(**x**_*k*_ − **x**_*n*_) = 1] and Var [𝕀δ*A*(**x**_*k*_ − **x**_*n*_)] = ℙ [𝕀δ*A*(**x**_*k*_ − **x**_*n*_) = 1] − (ℙ [𝕀δ*A*(**x**_*k*_ − **x**_*n*_) = 1])^2^ [52], we rewrite Equation (E.1) as,

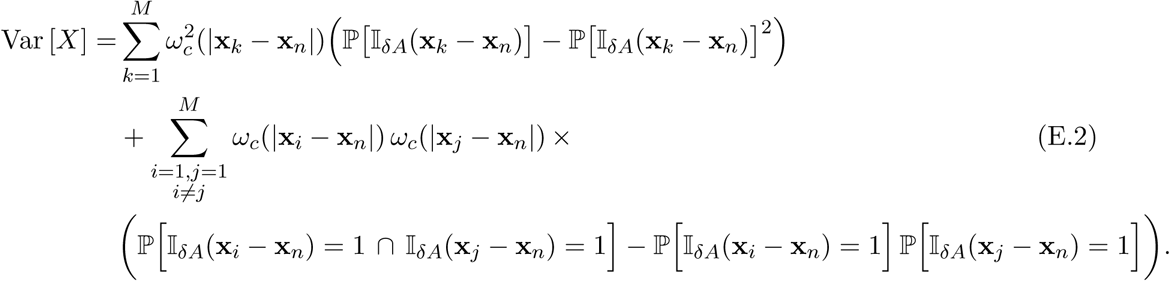

Using Equation (D.4), we write,

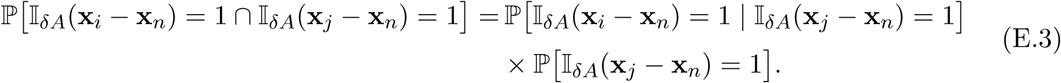

These definitions allow us to rewrite Equation (E.2) in terms of continuous variables by multiplying the corresponding conditional probabilities for the presence of individuals with the interaction kernels and summing over all possible displacements as,

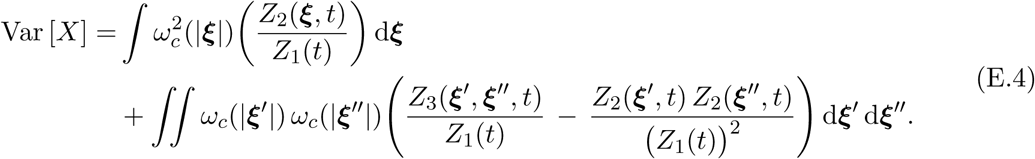

### F Comparison of moment closure methods

The moment closure methods help to approximate the third-order spatial moments in terms of the first and second spatial moments. In this section, we compare the performance of popular moment closure methods. The closure methods considered are the power-1 closure (P1), the symmetric power-2 closure (P2S), the asymmetric power-2 closure (P2A) and the Kirkwood superposition approximation (KSA) [29, 53].

The power-1 closure (P1) method use an approximation for the third moment, *Z*_3_(***ξ, ξ***′, *t*), given by,

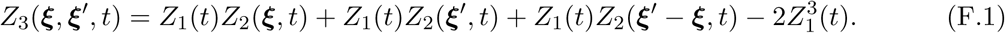

The symmetric power-2 closure is given by,

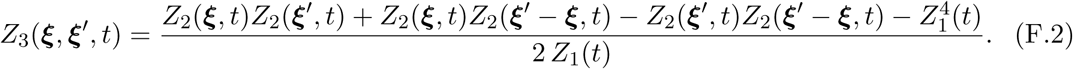

The asymmetric power-2 closure is given by,

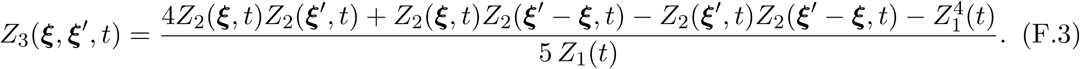

The Kirkwood superposition approximation (KSA) is given by,

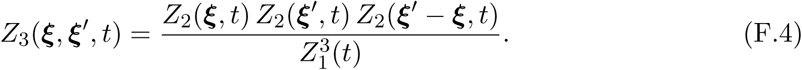

To compare the accuracy of closure methods, we compute the solutions of the spatial moment model with each of the four closure methods and compare it with the averaged data from the IBM simulation. We calculate the density dynamics and pair-correlation function for a population with initial population size, *N* (0) = 150, and random initial arrangement of individuals. The density dynamics computed using all four closure methods, as well as from the IBM simulations for this population, are shown in Figure 10(a). Our results indicate that the asymmetric power-2 closure provides the best match with the average results from the IBM. Similarly, the pair-correlation function computed using the asymmetric power-2 closure most accurately reproduces *C*(|***ξ***|, *t*) from the IBM, as shown in Figure 10(b). While our results suggest that the asymmetric power-2 closure provides the best approximation to IBM for the parameters considered, the motive of this study is not to advocate for one particular closure approximation over others. Instead, we provide a general moment dynamics model that can be implemented using different closure methods if one particular closure approximation is preferred over others.

**Figure 10:**
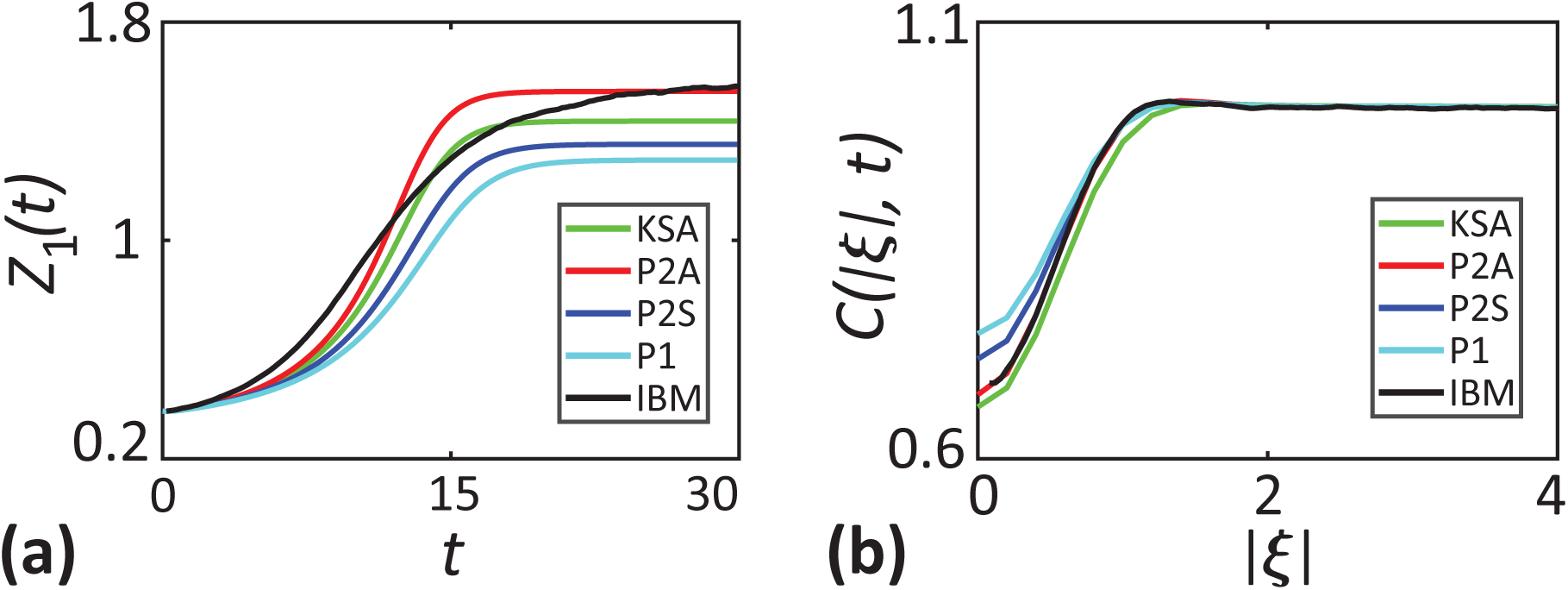
Comparison of moment closure methods. **a** shows the density of individuals as a function of time. **b** shows the *C*(|***ξ***|, *t*) computed at *t* = 30 as a function of separation distance. Parameter values are *σ*_*c*_ = 0.5, *γ*_*c*_ = 0.448, *σ*_*p*_ = *σ*_*d*_ = 4.0, *γ*_*p*_ = 0.009, *d* = 0.4, *p* = 0.2, *m* = 0.1, *µ*_*s*_ = 0.4 and *σ*_*s*_ = 0.1.

### G Numerical methods for solving the moment dynamics equation

Here we describe the numerical methods used for solving the dynamical equation for the second spatial moment, Equation (3.13). Temporal derivatives are approximated using an explicit Euler approximation implemented in MATLAB. The codes used are available on Github. The numerical scheme involves spatial discretisation of the displacement, ***ξ*** = (*ξ*_*x*_, *ξ*_*y*_), over the domain {−*ξ*_max_ ≤ *ξ*_*x*_, *ξ*_*y*_ ≤ *ξ*_max_} using a constant grid spacing of Δ***ξ***. We use a sufficiently large *ξ*_max_ such that 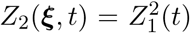 at the boundary since we anticipate that the usual mean-field condition will hold for sufficiently large displacements. We approximate the integral terms in Equations (3.11)-(3.13) using the trapezoid rule. The evaluation of these integral terms require the values of *Z*_2_(***ξ*** + ***ξ***′, *t*) for all values of ***ξ*** and ***ξ***′. A potential issue here is when both ***ξ*** and ***ξ***′ are large, there is a possibility that *Z*_2_(***ξ*** + ***ξ***′, *t*) lies outside of the computational domain. In such cases, we replace those terms with the value of *Z*_2_(***ξ***, *t*) at the boundary, *Z*_2_((*ξ*_max_, *ξ*_max_), *t*). The movement and dispersal kernels are normalised such that ***∫**µ*_*m*_(***ξ***) d***ξ*** = 1 and ***∫**µ*_*p*_(***ξ***) d***ξ*** = 1, using the trapezoid rule.

Solving for the dynamics of the second spatial moment, Equation (3.13), requires the evaluation of *Z*_1_(*t*). Since we consider a sufficiently large computational domain compared to the interaction ranges, the usual mean-field condition, 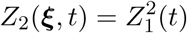 will be valid at large displacements. Using this property, we evaluate the first moments without actually solving the Equation (3.10). At each time step, the first moment is computed using, 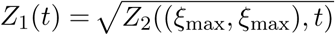. To compare the results from the spatial moment model with that of the IBM, we calculate the pair-correlation function as, 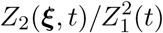. We use an initial condition, 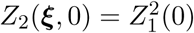. In all of our computation we use a constant time step, d*t* = 0.1, grid spacing, Δ***ξ*** = 0.2 and ***ξ***_max_ = 16. We find that these values of d*t*, and Δ***ξ*** are sufficiently small to produce grid-independent results. Further, we find that choosing larger values of ***ξ***_max_ does not affect our results.

### H Effect of short-range dispersal

Here, we investigate the effect of short-range dispersal of offspring in Figure 11. For these suites of simulations, we consider long-range competition and cooperation among individuals (*σ*_*c*_ = *σ*_*p*_ = 4.0) so that we can describe solely the dynamics resulting from the close dispersal of offspring (*σ*_*d*_ = 0.5). Again we consider three cases with initial population size, *N* (0) = 80, 240 and 400, where individuals are randomly distributed over the domain, as shown in Figure 11(a)-(c). The three initial conditions considered here are the same as those considered in Figure 4.

**Figure 11:**
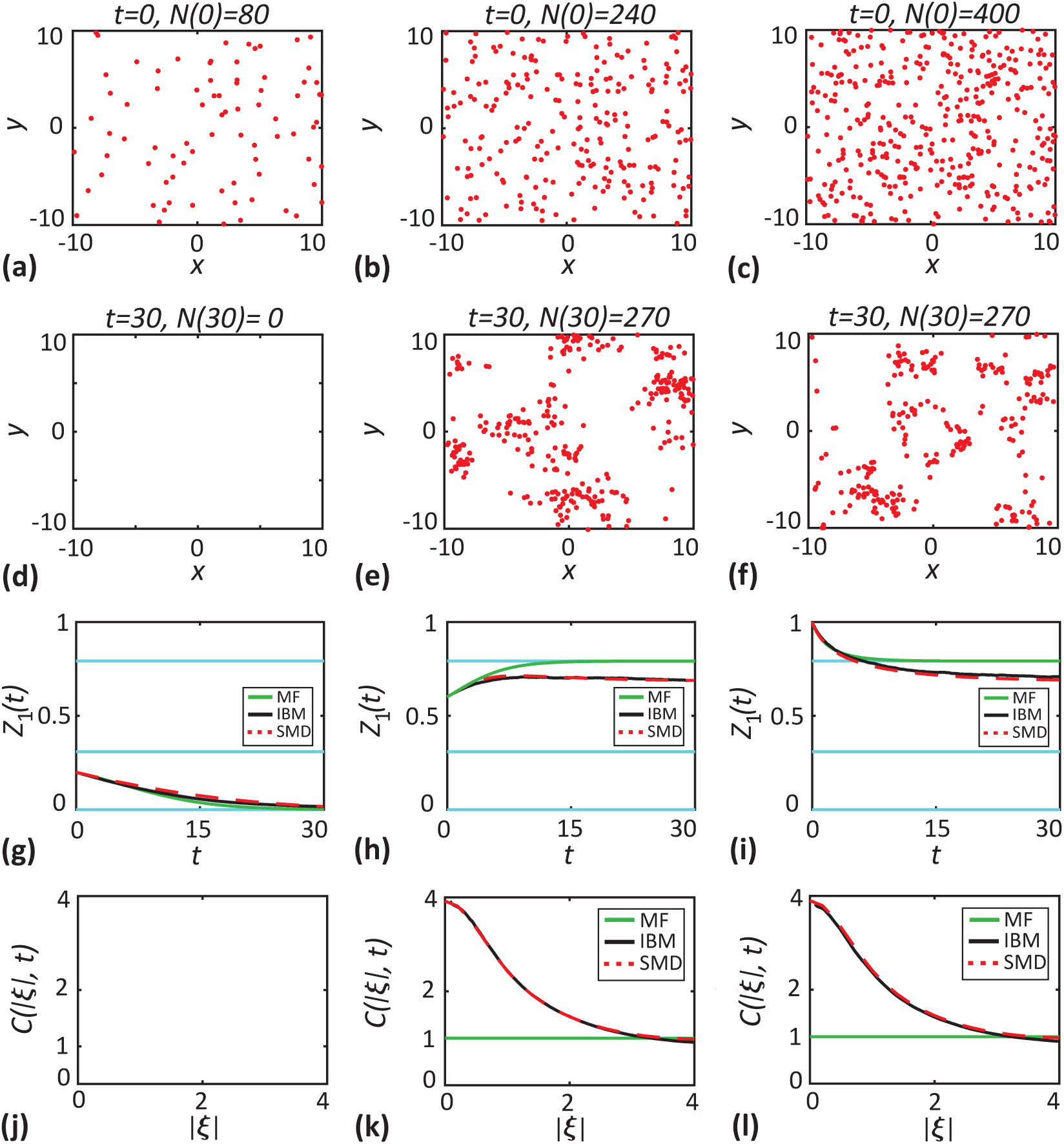
Effect of short-range dispersal. In these set of simulations, dispersal range is lowered to *σ*_*d*_ = 0.5. **a-c** show the initial locations of individuals (red dots) for three different population sizes, *N* (0) = 80, 240 and 400. **d-f** show the location of individuals at *t* = 30. **g-i** show the density of individuals as a function of time. Black solid lines correspond to the averaged results from 1000 realisations of the IBM, red dashed lines correspond to the solutions of spatial moment dynamics and green solid lines correspond to the solution of the mean-field model. The cyan lines show the critical densities. **j-l** show the *C*(|***ξ***|, *t*) computed at *t* = 30 as a function of separation distance. Parameter values are *d* = 0.4, *p* = 0.2, *m* = 0.1, *µ*_*s*_ = 0.4 and *σ*_*s*_ = 0.1.

When 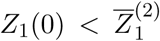, the population goes extinct since the total death rate is higher compared to the proliferation rate. For the two cases where 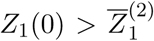, the population survives, and the locations of individuals at the final time are shown in Figure 11(e)-(f). As time progresses, individuals tend to be found in close groups corresponding to a clustered spatial structure. The clustering arises in the population through short-range dispersal, where offspring are placed in the close vicinity of parents. As time evolves, more and more individuals are placed in close neighbourhoods creating a strong clustering. Since we consider long-range competition, there is no mechanism to counteract or reduce the magnitude of the cluster formation. The presence of spatial clustering in the population is further confirmed by *C*(|***ξ***|, *t*) > 1 in Figure 11(k)-(l). Now the competition within these clusters enhances the net death rate. As a result, the population grows to a carrying capacity that is lower than 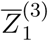. The spatial moment model accurately captures cluster formation due to short-scale dispersal and the resulting reduction in the carrying capacity.

