## Supplementary Material for "Population dynamics with spatial structure and an Allee effect"

#### Contents

**1 Comparison of long-range and short-range competition** **2**

**2 Numerical implementation of the individual-based model** **4**

**3 Definition of spatial moments** **6**

**4 Conditional probabilities for the presence of individuals** **7**

**5 Computation of variance** **9**

**6 Comparison of moment closure methods** **10**

**7 Numerical methods for solving the moment dynamics equation** **12**

**8 Effect of short-range dispersal** **13**

### 1 Comparison of long-range and short-range competition

Here we present a second set of results comparing the impact of short-range and long-range competition on the death rate, $\mathcal{D}_n$, analogous to the results in Figure 1. These additional results, in Figure S1, are generated with precisely the same parameters and spatial arrangement of individuals as those in Figure 1, with the exception that here we consider a quadratic functional form for the death rate ($F(X_n) = X_n^2$). Comparing the results in Figure 1 with the additional results in Figure S1 indicates that the choice of $F(X_n) = X_n^2$, reduces the death rates of individuals.

Similar to the results in Figure 1, the long-range competition leads to $\mathcal{D}_n$ being influenced by neighbours that are further apart, as shown in Figure S1(a)-(b). But the difference here is that overall we see a decrease in the death rate. For example, even though the death rate of the relatively isolated individual shown with the green dot is non-zero due to the long-range competition, the value of $\mathcal{D}_n = 0.076$ here is lower than $\mathcal{D}_n = 0.275$ computed using $F(X_n) = X_n$ in Figure 1. Under short-range competition, as shown in Figure S1(c)-(d), only the contribution from immediate neighbours is significant. As a result, the isolated individual (green dot), does not experience competition from its neighbours, leading to $\mathcal{D}_n = 0$.

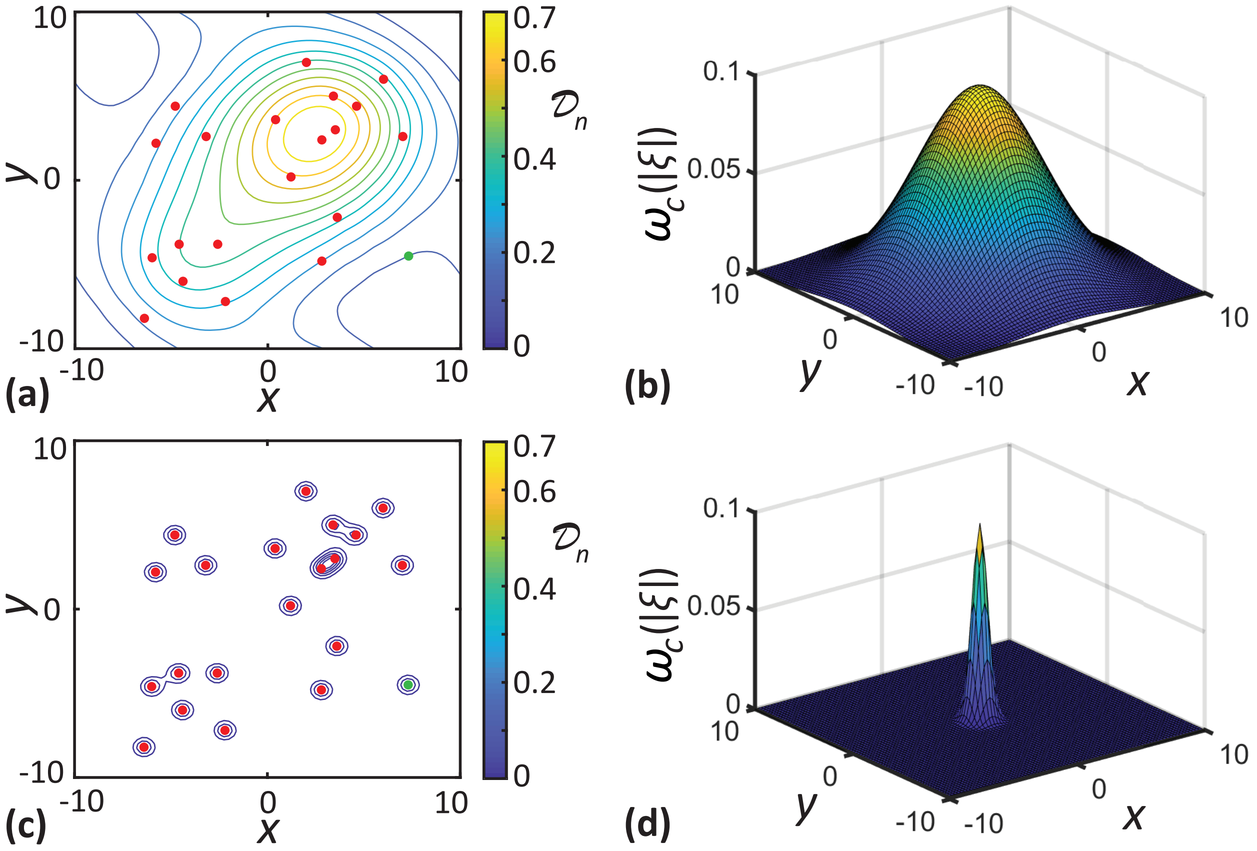


Figure S1: Visualisation of the impact of long-range and short-range neighbour-dependent interactions. Results in **a, c** show the location of individuals (red dots) superimposed with the level curves of $\mathcal{D}_n$ for long-range and short range-competition, respectively. Results in **b, d** show the long-range ($\sigma_c = 4.0$) and short-range ($\sigma_c = 0.5$) competition kernel, respectively, where these kernels are centred at the origin. For the computation of $\mathcal{D}_n$, we use $F(X_n) = X_n^2$ and $\gamma_c = 0.1$.

### 2 Numerical implementation of the individual-based model

Here we outline the numerical implementation of our individual-based model (IBM) using the Gillespie algorithm [1]. The codes used for the simulation of the IBM are available on Github. In each simulation, we initially populate the computational domain, of size $L \times L$, with $N(0)$ individuals placed at random. We use periodic boundary conditions along all boundaries. For each potential event, we compute the death and proliferation rates of individuals using Equation (2.3) and Equation (2.6). The movement of individuals is density independent, and the constant intrinsic movement rate is $m$. The sum of event rates of all individuals is given by,

$$\lambda(t) = \sum_{n=1}^{N(t)} \left( \mathcal{D}_n + \mathcal{P}_n + m \right). \tag{S1}$$

Each time the IBM is updated, one of the three possible events occurs, and the time interval between successive events is exponentially distributed with mean $1/\lambda(t)$. The probability for any of the events to occur is proportional to the rate of the corresponding event. For a proliferation event, an offspring is placed at a displacement sampled from the dispersal kernel, $\mu_p(\boldsymbol{\xi})$, and the population size increases by one. For a death event, the population size reduces by one. For a movement event, an individual traverses a displacement that is sampled from the movement displacement kernel, $\mu_m(\boldsymbol{\xi})$.

We compute the average density of individuals at a particular time by dividing the population size, $N(t)$, by the area of the computational domain, $L^2$. To compute the pair-correlation function, $C(|\boldsymbol{\xi}|, t)$, we consider a reference individual located at $\mathbf{x}_n$ and calculate all distances $|\boldsymbol{\xi}| = |\mathbf{x}_k - \mathbf{x}_n|$, associated with the other $N(t) - 1$ individuals [2, 3]. We repeat this process with each of the remaining individuals until every individual has acted as the reference individual. The pair-correlation function is calculated by enumerating the distances which fall into the interval, $[|\boldsymbol{\xi}| - \delta|\boldsymbol{\xi}|/2, |\boldsymbol{\xi}| + \delta|\boldsymbol{\xi}|/2]$. We normalise the bin count by a factor of $2\pi|\boldsymbol{\xi}|\delta|\boldsymbol{\xi}|N(t)(N(t) - 1)/L^2$ to ensure that $C(|\boldsymbol{\xi}|, t) = 1$ in the absence of spatial structure. The choice of bin width, $\delta|\boldsymbol{\xi}|$, is crucial in computing the pair-correlation function. When $\delta|\boldsymbol{\xi}|$ is very small, we obtain a noise dominated $C(|\boldsymbol{\xi}|, t)$. In contrast, very large $\delta|\boldsymbol{\xi}|$ leads to an overly smooth $C(|\boldsymbol{\xi}|, t)$ that fails to describe the effects of short-range interactions. In all our simulations, we use an intermediate value of $\delta|\boldsymbol{\xi}| = 0.2$ which helps us to avoid the two extremities.

### 3 Definition of spatial moments

Here, we provide a more formal mathematical definition for the spatial moments [4–6]. Let us suppose $D_{\delta A}(\mathbf{x}) \subset \mathbb{R}^2$ is a disc of area $\delta A$ centred at position $\mathbf{x} \in \mathbb{R}^2$ and the number of individuals in the region $D_{\delta A}(\mathbf{x})$, at a time $t$, is denoted by the random variable $N(D_{\delta A}(\mathbf{x}), t)$. The first spatial moment, $Z_1(t)$, can be computed by dividing the population size of individuals by the area of the domain. Hence we have,

$$Z_1(t) = \lim_{\delta A \to 0} \frac{1}{\delta A} \mathbb{E}\Big[N\big(D_{\delta A}(\mathbf{x}), t\big)\Big]. \tag{S2}$$

The second spatial moment, $Z_2(\boldsymbol{\xi}, t)$, is the average density of pairs of individuals separated by a displacement $\boldsymbol{\xi}$ at time $t$. For a pair of individuals separated by a displacement $\boldsymbol{\xi}$, we have,

$$Z_2(\boldsymbol{\xi}, t) = \lim_{\delta A \to 0} \frac{1}{\delta A^2} \mathbb{E}\Big[N\left(D_{\delta A}(\mathbf{x}), t\right) N\left(D_{\delta A}(\mathbf{x}+\boldsymbol{\xi}), t\right) - N\left(D_{\delta A}(\mathbf{x}) \cap D_{\delta A}(\mathbf{x}+\boldsymbol{\xi}), t\right)\Big]. \tag{S3}$$

The second term in the expectation in Equation (S3) is necessary to avoid counting self-pairs. For non-overlapping regions $D_{\delta A}(\mathbf{x})$ and $D_{\delta A}(\mathbf{x}+\boldsymbol{\xi})$, this term becomes zero as $\delta A \to 0$. The third spatial moment is the density of triplets of individuals, and is similarly defined as,

$$\begin{aligned} Z_3(\boldsymbol{\xi}, \boldsymbol{\xi}', t) = \lim_{\delta A \to 0} \frac{1}{\delta A^3} \mathbb{E}\Big[ & N(D_{\delta A}(\mathbf{x}), t)\, N(D_{\delta A}(\mathbf{x}+\boldsymbol{\xi}), t)\, N(D_{\delta A}(\mathbf{x}+\boldsymbol{\xi}'), t) \\ & - N(D_{\delta A}(\mathbf{x}) \cap D_{\delta A}(\mathbf{x}+\boldsymbol{\xi}), t)\, N(D_{\delta A}(\mathbf{x}+\boldsymbol{\xi}'), t) \\ & - N(D_{\delta A}(\mathbf{x}) \cap D_{\delta A}(\mathbf{x}+\boldsymbol{\xi}'), t)\, N(D_{\delta A}(\mathbf{x}+\boldsymbol{\xi}), t) \\ & - N(D_{\delta A}(\mathbf{x}+\boldsymbol{\xi}) \cap D_{\delta A}(\mathbf{x}+\boldsymbol{\xi}'), t)\, N(D_{\delta A}(\mathbf{x}), t) \\ & + 2N(D_{\delta A}(\mathbf{x}) \cap D_{\delta A}(\mathbf{x}+\boldsymbol{\xi}') \cap D_{\delta A}(\mathbf{x}+\boldsymbol{\xi}), t)\Big]. \end{aligned} \tag{S4}$$

Again, the extra terms in Equation (S4) are needed to avoid counting non-distinct triplets.

### 4 Conditional probabilities for the presence of individuals

In this section we derive expressions for the probabilities of finding individuals at specific displacements conditional on the presence of other individuals. In the limit, $\delta A \to 0$, the probability of having one individual in the region $D_{\delta A}(\mathbf{x})$ is given by,

$$\mathbb{P}\left[N\big(D_{\delta A}(\mathbf{x}),t\big)=1\right]=\mathbb{E}\Big[N\big(D_{\delta A}(\mathbf{x}),t\big)\Big]. \tag{S5}$$

Now, the probability of having two individuals located in non overlapping regions $D_{\delta A}(\mathbf{x})$ and $D_{\delta A}(\mathbf{x}+\boldsymbol{\xi})$, respectively, is given by,

$$\mathbb{P}\Big[N\big(D_{\delta A}(\mathbf{x}),t\big)=1 \cap N\big(D_{\delta A}(\mathbf{x}+\boldsymbol{\xi}),t\big)=1\Big]=\mathbb{E}\Big[N\big(D_{\delta A}(\mathbf{x}),t\big)\,N\big(D_{\delta A}(\mathbf{x}+\boldsymbol{\xi}),t\big)\Big]. \tag{S6}$$

Similarly, the probability for having three individuals at non overlapping regions $D_{\delta A}(\mathbf{x})$, $D_{\delta A}(\mathbf{x}+\boldsymbol{\xi})$ and $D_{\delta A}(\mathbf{x}+\boldsymbol{\xi}')$, respectively, is given by,

$$\begin{aligned}\mathbb{P}\Big[N\big(D_{\delta A}(\mathbf{x}),t\big)=1 \cap N\big(D_{\delta A}(\mathbf{x}+\boldsymbol{\xi}),t\big)=1 \cap N\big(D_{\delta A}(\mathbf{x}+\boldsymbol{\xi}'),t\big)=1\Big]\\ =\mathbb{E}\Big[N\big(D_{\delta A}(\mathbf{x}),t\big)\,N\big(D_{\delta A}(\mathbf{x}+\boldsymbol{\xi}),t\big)\,N\big(D_{\delta A}(\mathbf{x}+\boldsymbol{\xi}'),t\big)\Big].\end{aligned} \tag{S7}$$

To compute the event rates, we need to find the probabilities of individuals being present at a given displacement, conditional on the presence of other individuals. To compute the conditional probabilities, we use the property that,

$$\mathbb{P}[A \mid B]=\frac{\mathbb{P}[A\cap B]}{\mathbb{P}[B]}. \tag{S8}$$

The conditional probability of finding an individual at a displacement $\mathbf{x}+\boldsymbol{\xi}$, given that the reference individual is located at $\mathbf{x}$, is,

$$\mathbb{P}\Big[N(D_{\delta A}(\mathbf{x}+\boldsymbol{\xi}),t)=1 \mid N(D_{\delta A}(\mathbf{x}),t)=1\Big]=\frac{\mathbb{P}\Big[N(D_{\delta A}(\mathbf{x}+\boldsymbol{\xi}),t)=1\cap N(D_{\delta A}(\mathbf{x}),t)=1\Big]}{\mathbb{P}\Big[N(D_{\delta A}(\mathbf{x}),t)=1\Big]}. \tag{S9}$$

Using the definitions of probabilities in Equations (S5)-(S6) and the definitions of spatial moments in Equations (S2)-(S3), we rewrite the the numerator and denominator of Equation

(S9) as,

$$\mathbb{P}\Big[N(D_{\delta A}(\mathbf{x}),t)=1\Big]=Z_1(t)\,\delta A, \tag{S10}$$

$$\mathbb{P}\Big[N(D_{\delta A}(\mathbf{x}+\boldsymbol{\xi}),t)=1\cap N(D_{\delta A}(\mathbf{x}),t)=1\Big]=Z_2(\boldsymbol{\xi},t)\,(\delta A)^2. \tag{S11}$$

Hence the conditional probability of finding an individual at a displacement $\mathbf{x}+\boldsymbol{\xi}$ from a reference individual at $\mathbf{x}$ is given by,

$$\mathbb{P}\Big[N(D_{\delta A}(\mathbf{x}+\boldsymbol{\xi}),t)=1 \mid N(D_{\delta A}(\mathbf{x}),t)=1\Big]=\frac{Z_2(\boldsymbol{\xi},t)\,\delta A}{Z_1(t)}. \tag{S12}$$

Similarly, we compute the conditional probability of finding an individual at a displacement $\mathbf{x}+\boldsymbol{\xi'}$, given that a pair of individuals where constituent individuals are located at $\mathbf{x}+\boldsymbol{\xi}$ and $\mathbf{x}$, respectively, as

$$\begin{aligned}
&\mathbb{P}\Big[N(D_{\delta A}(\mathbf{x}+\boldsymbol{\xi'}),t)=1 \mid N(D_{\delta A}(\mathbf{x}+\boldsymbol{\xi}),t)=1\cap N(D_{\delta A}(\mathbf{x}),t)=1\Big]\\
&=\frac{\mathbb{P}\Big[N(D_{\delta A}(\mathbf{x}+\boldsymbol{\xi'}),t)=1\cap N(D_{\delta A}(\mathbf{x}+\boldsymbol{\xi}),t)=1\cap N(D_{\delta A}(\mathbf{x}),t)=1\Big]}{\mathbb{P}\Big[N(D_{\delta A}(\mathbf{x}+\boldsymbol{\xi}),t)=1\cap N(D_{\delta A}(\mathbf{x}),t)=1\Big]},\\
&=\frac{Z_3(\boldsymbol{\xi},\boldsymbol{\xi'},t)\,\delta A}{Z_2(\boldsymbol{\xi},t)}.
\end{aligned} \tag{S13}$$

### 5 Computation of variance

Here we outline the steps involved in the derivation of expression for variance of the neighbourhood density in Equation (3.7). The fundamental definition of the variance is,

$$
\begin{aligned}
\mathrm{Var}\,[X] =& \mathrm{Var}\left[\sum_{k=1}^{M} \omega_c(|\mathbf{x}_k - \mathbf{x}_n|)\, \mathbb{I}_{\delta A}(\mathbf{x}_k - \mathbf{x}_n)\right], \\
=& \sum_{k=1}^{M} \omega_c^2(|\mathbf{x}_k - \mathbf{x}_n|)\, \mathrm{Var}\Big[\mathbb{I}_{\delta A}(\mathbf{x}_k - \mathbf{x}_n)\Big] \\
&+ \sum_{\substack{i=1,j=1\\ i\neq j}}^{M} \omega_c(|\mathbf{x}_i - \mathbf{x}_n|)\, \omega_c(|\mathbf{x}_j - \mathbf{x}_n|)\, \mathrm{Cov}\left[\mathbb{I}_{\delta A}(\mathbf{x}_i - \mathbf{x}_n),\ \mathbb{I}_{\delta A}(\mathbf{x}_j - \mathbf{x}_n)\right].
\end{aligned}
\tag{S14}
$$

Now, using the properties of indicator function, $\mathbb{E}\big[\mathbb{I}_{\delta A}(\mathbf{x}_k - \mathbf{x}_n)\big] = \mathbb{P}\big[\mathbb{I}_{\delta A}(\mathbf{x}_k - \mathbf{x}_n) = 1\big]$ and $\mathrm{Var}\big[\mathbb{I}_{\delta A}(\mathbf{x}_k - \mathbf{x}_n)\big] = \mathbb{P}\big[\mathbb{I}_{\delta A}(\mathbf{x}_k - \mathbf{x}_n) = 1\big] - \Big(\mathbb{P}\big[\mathbb{I}_{\delta A}(\mathbf{x}_k - \mathbf{x}_n) = 1\big]\Big)^2$ [7], we rewrite Equation (S14) as,

$$
\begin{aligned}
\mathrm{Var}\,[X] =& \sum_{k=1}^{M} \omega_c^2(|\mathbf{x}_k - \mathbf{x}_n|)\Big(\mathbb{P}\big[\mathbb{I}_{\delta A}(\mathbf{x}_k - \mathbf{x}_n)\big] - \mathbb{P}\big[\mathbb{I}_{\delta A}(\mathbf{x}_k - \mathbf{x}_n)\big]^2\Big) \\
&+ \sum_{\substack{i=1,j=1\\ i\neq j}}^{M} \omega_c(|\mathbf{x}_i - \mathbf{x}_n|)\, \omega_c(|\mathbf{x}_j - \mathbf{x}_n|) \times \\
&\Bigg(\mathbb{P}\Big[\mathbb{I}_{\delta A}(\mathbf{x}_i - \mathbf{x}_n) = 1 \cap \mathbb{I}_{\delta A}(\mathbf{x}_j - \mathbf{x}_n) = 1\Big] - \mathbb{P}\Big[\mathbb{I}_{\delta A}(\mathbf{x}_i - \mathbf{x}_n) = 1\Big]\, \mathbb{P}\Big[\mathbb{I}_{\delta A}(\mathbf{x}_j - \mathbf{x}_n) = 1\Big]\Bigg).
\end{aligned}
\tag{S15}
$$

Using Equation (S8), we write,

$$
\begin{aligned}
\mathbb{P}\big[\mathbb{I}_{\delta A}(\mathbf{x}_i - \mathbf{x}_n) = 1 \cap \mathbb{I}_{\delta A}(\mathbf{x}_j - \mathbf{x}_n) = 1\big] =& \mathbb{P}\big[\mathbb{I}_{\delta A}(\mathbf{x}_i - \mathbf{x}_n) = 1 \mid \mathbb{I}_{\delta A}(\mathbf{x}_j - \mathbf{x}_n) = 1\big] \\
&\times \mathbb{P}\big[\mathbb{I}_{\delta A}(\mathbf{x}_j - \mathbf{x}_n) = 1\big].
\end{aligned}
\tag{S16}
$$

These definitions allow us to rewrite Equation (S15) in terms of continuous variables by multiplying the corresponding conditional probabilities for the presence of individuals with the interaction kernels and summing over all possible displacements as,

$$
\begin{aligned}
\mathrm{Var}\,[X] =& \int \omega_c^2(|\boldsymbol{\xi}|)\left(\frac{Z_2(\boldsymbol{\xi}, t)}{Z_1(t)}\right) \mathrm{d}\boldsymbol{\xi} \\
&+ \iint \omega_c(|\boldsymbol{\xi}'|)\, \omega_c(|\boldsymbol{\xi}''|)\left(\frac{Z_3(\boldsymbol{\xi}', \boldsymbol{\xi}'', t)}{Z_1(t)} - \frac{Z_2(\boldsymbol{\xi}', t)\, Z_2(\boldsymbol{\xi}'', t)}{\big(Z_1(t)\big)^2}\right) \mathrm{d}\boldsymbol{\xi}'\, \mathrm{d}\boldsymbol{\xi}''.
\end{aligned}
\tag{S17}
$$

### 6 Comparison of moment closure methods

The moment closure methods help to approximate the third-order spatial moments in terms of the first and second spatial moments. In this section, we compare the performance of popular moment closure methods. The closure methods considered are the power-1 closure (P1), the symmetric power-2 closure (P2S), the asymmetric power-2 closure (P2A) and the Kirkwood superposition approximation (KSA) [2,8].

The power-1 closure (P1) method use an approximation for the third moment, $Z_3(\boldsymbol{\xi}, \boldsymbol{\xi}', t)$, given by,

$$Z_3(\boldsymbol{\xi}, \boldsymbol{\xi}', t) = Z_1(t)Z_2(\boldsymbol{\xi}, t) + Z_1(t)Z_2(\boldsymbol{\xi}', t) + Z_1(t)Z_2(\boldsymbol{\xi}' - \boldsymbol{\xi}, t) - 2Z_1^3(t). \tag{S18}$$

The symmetric power-2 closure is given by,

$$Z_3(\boldsymbol{\xi}, \boldsymbol{\xi}', t) = \frac{Z_2(\boldsymbol{\xi}, t)Z_2(\boldsymbol{\xi}', t) + Z_2(\boldsymbol{\xi}, t)Z_2(\boldsymbol{\xi}' - \boldsymbol{\xi}, t) - Z_2(\boldsymbol{\xi}', t)Z_2(\boldsymbol{\xi}' - \boldsymbol{\xi}, t) - Z_1^4(t)}{2\, Z_1(t)}. \tag{S19}$$

The asymmetric power-2 closure is given by,

$$Z_3(\boldsymbol{\xi}, \boldsymbol{\xi}', t) = \frac{4Z_2(\boldsymbol{\xi}, t)Z_2(\boldsymbol{\xi}', t) + Z_2(\boldsymbol{\xi}, t)Z_2(\boldsymbol{\xi}' - \boldsymbol{\xi}, t) - Z_2(\boldsymbol{\xi}', t)Z_2(\boldsymbol{\xi}' - \boldsymbol{\xi}, t) - Z_1^4(t)}{5\, Z_1(t)}. \tag{S20}$$

The Kirkwood superposition approximation (KSA) is given by,

$$Z_3(\boldsymbol{\xi}, \boldsymbol{\xi}', t) = \frac{Z_2(\boldsymbol{\xi}, t)\, Z_2(\boldsymbol{\xi}', t)\, Z_2(\boldsymbol{\xi}' - \boldsymbol{\xi}, t)}{Z_1^3(t)}. \tag{S21}$$

To compare the accuracy of closure methods, we compute the solutions of the spatial moment model with each of the four closure methods and compare it with the averaged data from the IBM simulation. We calculate the density dynamics and pair-correlation function for a population with initial population size, $N(0) = 150$, and random initial arrangement of individuals. The density dynamics computed using all four closure methods, as well as from the IBM simulations for this population, are shown in Figure S2(a). Our results indicate that the asymmetric power-2 closure provides the best match with the average results from the IBM. Similarly, the pair-correlation function computed using the asymmetric power-2 closure most accurately reproduces $C(|\boldsymbol{\xi}|, t)$ from the IBM, as shown in Figure S2(b). While our results suggest that the asymmetric power-2 closure provides the best approximation to IBM for the parameters considered, the motive of this study is not to advocate for one particular closure approximation over others. Instead, we provide a general moment dynamics model that can be implemented using different closure methods if one particular closure approximation is preferred over others.

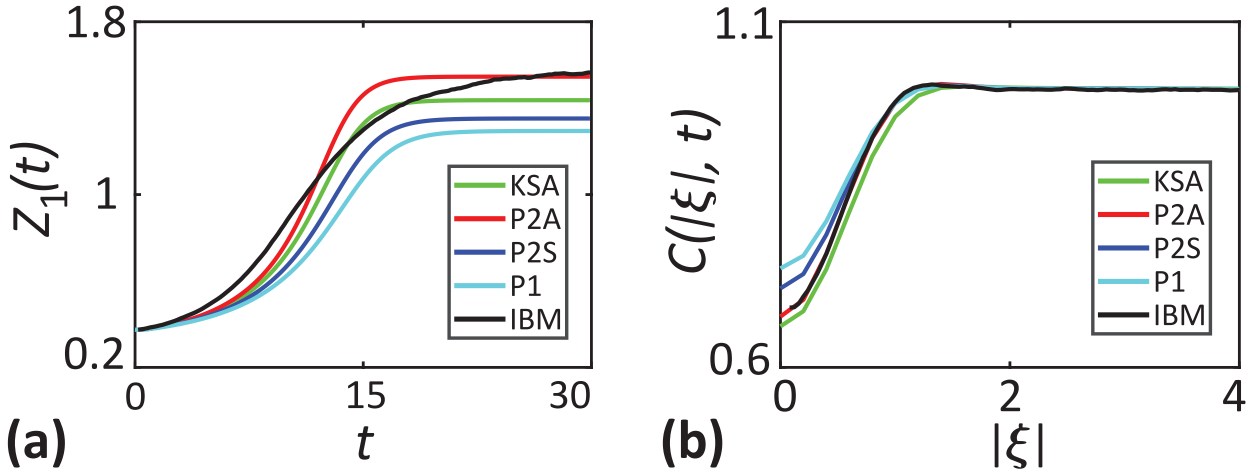


Figure S2: Comparison of moment closure methods. **a** shows the density of individuals as a function of time. **b** shows the $C(|\boldsymbol{\xi}|, t)$ computed at $t = 30$ as a function of separation distance. Parameter values are $\sigma_c = 0.5, \gamma_c = 0.448, \sigma_p = \sigma_d = 4.0, \gamma_p = 0.009, d = 0.4, p = 0.2, m = 0.1, \mu_s = 0.4$ and $\sigma_s = 0.1$.

### 7 Numerical methods for solving the moment dynamics equation

Here we describe the numerical methods used for solving the dynamical equation for the second spatial moment, Equation (3.13). Temporal derivatives are approximated using an explicit Euler approximation implemented in MATLAB. The codes used are available on Github. The numerical scheme involves spatial discretisation of the displacement, $\boldsymbol{\xi} = (\xi_x, \xi_y)$, over the domain $\{-\xi_{\max} \leq \xi_x, \xi_y \leq \xi_{\max}\}$ using a constant grid spacing of $\Delta\boldsymbol{\xi}$. We use a sufficiently large $\xi_{\max}$ such that $Z_2(\boldsymbol{\xi}, t) = Z_1^2(t)$ at the boundary since we anticipate that the usual mean-field condition will hold for sufficiently large displacements. We approximate the integral terms in Equations (3.11)-(3.13) using the trapezoid rule. The evaluation of these integral terms require the values of $Z_2(\boldsymbol{\xi} + \boldsymbol{\xi}', t)$ for all values of $\boldsymbol{\xi}$ and $\boldsymbol{\xi}'$. A potential issue here is when both $\boldsymbol{\xi}$ and $\boldsymbol{\xi}'$, are large, there is a possibility that $Z_2(\boldsymbol{\xi}+\boldsymbol{\xi}', t)$ lies outside of the computational domain. In such cases, we replace those terms with the value of $Z_2(\boldsymbol{\xi}, t)$ at the boundary, $Z_2((\xi_{\max}, \xi_{\max}), t)$. The movement and dispersal kernels are normalised such that $\int \mu_m(\boldsymbol{\xi})\, \mathrm{d}\boldsymbol{\xi} = 1$ and $\int \mu_p(\boldsymbol{\xi})\, \mathrm{d}\boldsymbol{\xi} = 1$, using the trapezoid rule.

Solving for the dynamics of the second spatial moment, Equation (3.13), requires the evaluation of $Z_1(t)$. Since we consider a sufficiently large computational domain compared to the interaction ranges, the usual mean-field condition, $Z_2(\boldsymbol{\xi}, t) = Z_1^2(t)$ will be valid at large displacements. Using this property, we evaluate the first moments without actually solving the Equation (3.10). At each time step, the first moment is computed using, $Z_1(t) = \sqrt{Z_2((\xi_{\max}, \xi_{\max}), t)}$. To compare the results from the spatial moment model with that of the IBM, we calculate the pair-correlation function as, $Z_2(\boldsymbol{\xi}, t)/Z_1^2(t)$. We use an initial condition, $Z_2(\boldsymbol{\xi}, 0) = Z_1^2(0)$. In all of our computation we use a constant time step, $\mathrm{d}t = 0.1$, grid spacing, $\Delta\boldsymbol{\xi} = 0.2$ and $\boldsymbol{\xi}_{\max} = 16$. We find that these values of $\mathrm{d}t$, and $\Delta\boldsymbol{\xi}$ are sufficiently small to produce grid-independent results. Further, we find that choosing larger values of $\boldsymbol{\xi}_{\max}$ does not affect our results.

### 8 Effect of short-range dispersal

Here, we investigate the effect of short-range dispersal of offspring in Figure S3. For these suites of simulations, we consider long-range competition and cooperation among individuals ($\sigma_c = \sigma_p = 4.0$) so that we can describe solely the dynamics resulting from the close dispersal of offspring ($\sigma_d = 0.5$). Again we consider three cases with initial population size, $N(0) = 80, 240$ and $400$, where individuals are randomly distributed over the domain, as shown in Figure S3(a)-(c). The three initial conditions considered here are the same as those considered in Figure 4.

When $Z_1(0) < \overline{Z}_1^{(2)}$, the population goes extinct since the total death rate is higher compared to the proliferation rate. For the two cases where $Z_1(0) > \overline{Z}_1^{(2)}$, the population survives, and the locations of individuals at the final time are shown in Figure S3(e)-(f). As time progresses, individuals tend to be found in close groups corresponding to a clustered spatial structure. The clustering arises in the population through short-range dispersal, where offspring are placed in the close vicinity of parents. As time evolves, more and more individuals are placed in close neighbourhoods creating a strong clustering. Since we consider long-range competition, there is no mechanism to counteract or reduce the magnitude of the cluster formation. The presence of spatial clustering in the population is further confirmed by $C(|\boldsymbol{\xi}|, t) > 1$ in Figure S3(k)-(l). Now the competition within these clusters enhances the net death rate. As a result, the population grows to a carrying capacity that is lower than $\overline{Z}_1^{(3)}$. The spatial moment model accurately captures cluster formation due to short-scale dispersal and the resulting reduction in the carrying capacity.

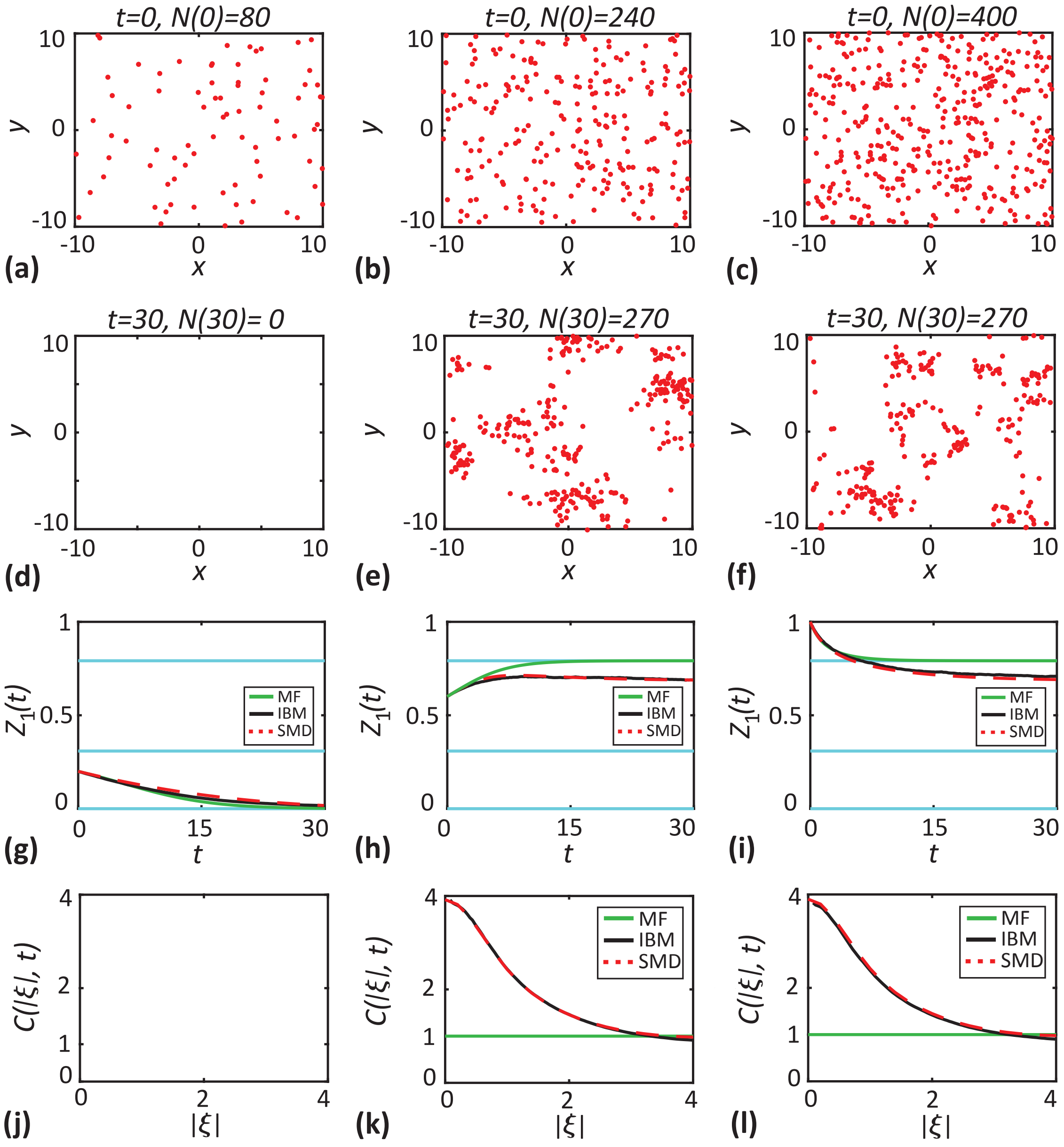


Figure S3: Effect of short-range dispersal. In these set of simulations, dispersal range is lowered to $\sigma_d = 0.5$. **a-c** show the initial locations of individuals (red dots) for three different population sizes, $N(0) = 80, 240$ and 400. **d-f** show the location of individuals at $t = 30$. **g-i** show the density of individuals as a function of time. Black solid lines correspond to the averaged results from 1000 realisations of the IBM, red dashed lines correspond to the solutions of spatial moment dynamics and green solid lines correspond to the solution of the mean-field model. The cyan lines show the critical densities. **j-l** show the $C(|\boldsymbol{\xi}|, t)$ computed at $t = 30$ as a function of separation distance. Parameter values are $d = 0.4, p = 0.2, m = 0.1, \mu_s = 0.4$ and $\sigma_s = 0.1$.
